# From the third to the seventh generation of selection for muscle fat content in rainbow trout: Consequences for flesh quality

**DOI:** 10.1101/2023.12.04.569019

**Authors:** Florence Lefevre, Jérôme Bugeon, Lionel Goardon, Thierry Kerneis, Laurent Labbe, Stéphane Panserat, Françoise Medale, Edwige Quillet

## Abstract

Muscle lipid content was shown to affect many quality features in salmonids in both raw and processed fillets. The objective of the present work was to assess the consequences of 7 generations of divergent selection for muscle adiposity on some rainbow trout flesh quality and muscle parameters. Fish from Lean (L) and Fat (F) lines had a similar body weight but L fish were longer and had consequently lower condition factor values. Carcass yield was not affected by selective breeding, but L fish had lower hepato- and gonado-somatic indexes, a bigger head, and lower fillet yield than F fish. A difference of more than 15 points in mean fat-meter^®^ values (genetic selection criteria) was measured between the two lines. Mean muscle lipid content was 5.0±1.0% for L line vs 13.5±2.2% for F line. An absolute difference of more than 6% was measured in fillet dry matter content between the two lines, for raw, cooked, and smoked fillets. Raw fillets from F fish were lighter (L*>) and more colorful (a* and b*>), but softer than those from the L line. Quality parameters of cooked fillets were very similar between the two lines, whereas smoked fillets exhibited, between the two lines, similar differences than raw fillets. A large difference in white muscle fiber size was observed, fish from F line having higher fiber mean diameter, a lower proportion of small fibers, and a higher proportion of large fibers. Sex effects were observed on these immature fish, on classically sex-related traits (GSI and head development), but also on muscle fiber size. Moreover, these effects were more marked in F line. Correlation analysis showed that raw fillet color was positively related to muscle adiposity whereas mechanical resistance was negatively related. Raw fillet mechanical resistance was also negatively correlated to white muscle fiber size. Moreover, smoked fillet quality parameters were correlated to raw fillet ones. The relationships between muscle adiposity, but also muscle cellularity, and fillet quality were discussed.

**Highlights:**

- 7 generations of adiposity divergent selection affect raw and smoked fillet quality.
- Selective breeding led to a noteworthy response in white muscle cellularity.
- Some differences between male and female was measured in immature pan-size trout.

**Graphical abstract:** 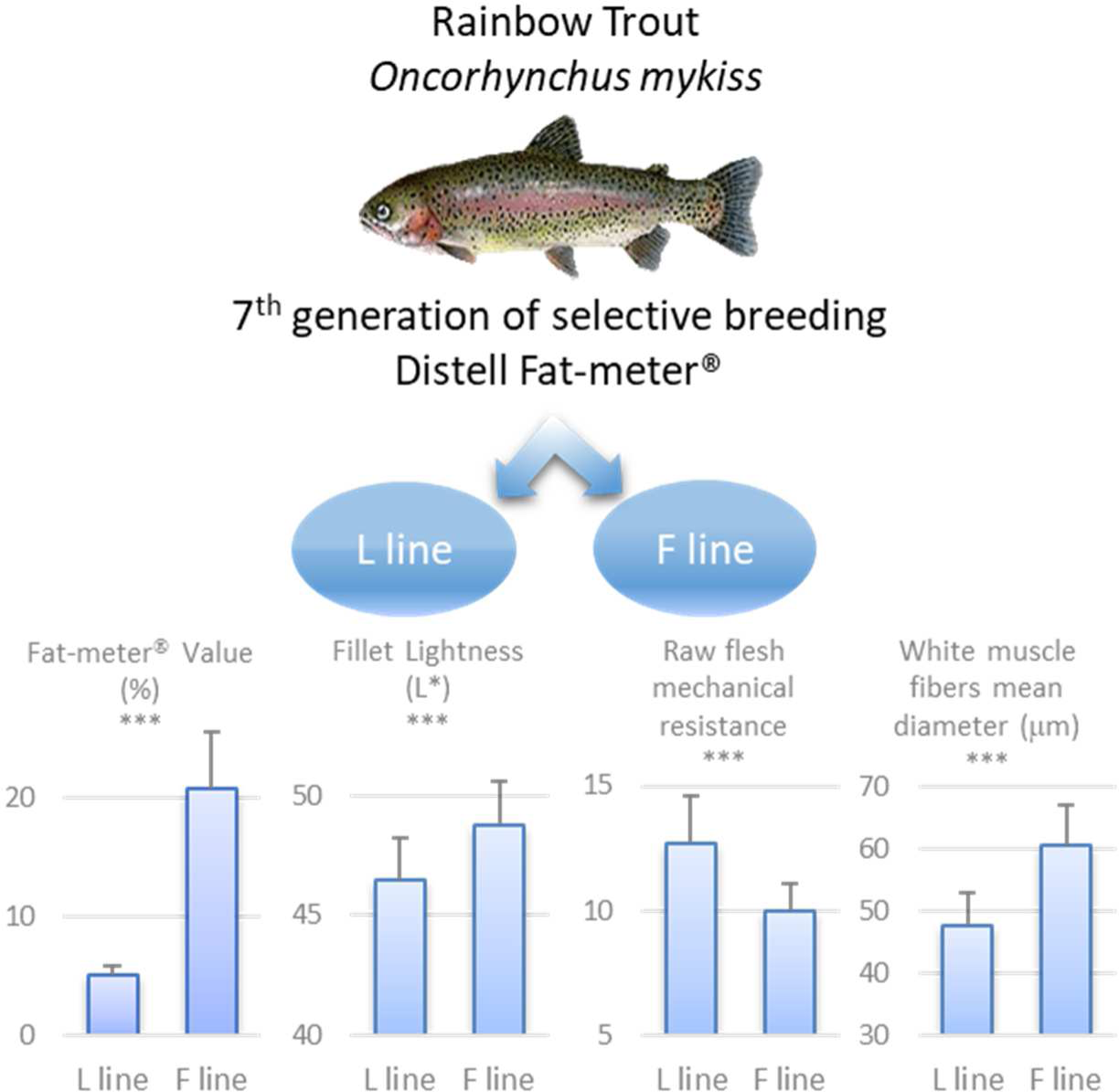

## 1. Introduction

In fish, as in other muscle food, lipid content was shown to determine nutritional quality, but also to affect organoleptic properties. Salmonid are an especially relevant model to study the relationship between muscle lipid content and flesh quality, as muscle lipid content can greatly vary from a few percent to over 10% in rainbow trout and even more than 15% in Atlantic salmon (Medale, 2009). Salmonid muscle adiposity depends on fish physiological stage (age, sexual maturation, for example), but rearing practices can control global and muscle fish adiposity whether by nutritional strategy or selective breeding. Effective nutritional strategy to increase fish adiposity are whether increasing feed ration, diet energy content or diet lipid content (Weil et al., 2013). Selective breeding was otherwise shown to be an effective way to control trout adiposity (Quillet et al., 2005; Tobin et al., 2006). In rainbow trout, a two ways divergent selection, using a non-destructive estimation of muscle lipid content (Distell Fish Fat Meter^®^), produced a lean (L) muscle line and a fat (F) muscle line (Quillet et al., 2005). After two generations of selection, large fish of the F line exhibited an increase by 15% to 31% in the mean muscle lipid content depending on the diet (Quillet et al., 2007). Further experiments with fish from the third generation of selection showed that the lines differed in fat allocation between visceral adipose tissue and muscle (Kolditz et al., 2008). Diploid pan-size trout from the same third generation of selection demonstrated an absolute difference of 5.5 % for fat-meter^®^ values, and of 2.5 % for muscle dry matter content (Lefevre et al., 2015).

Whatever the lever that affects muscle adiposity, muscle lipid content was shown to have consequences on flesh quality. Indeed, higher muscle fat content is generally associated to more colorful fillet, higher sensory scores for moisture or flavor, but somewhat less firm fillet (Lefevre et al., 2015; Morkore et al., 2001; Robb et al., 2002; Suarez et al., 2014). Those effects are globally similar for both nutritionally or genetically control of flesh adiposity. However, controlling muscle adiposity by genetic way allows estimating the relationship between muscle lipid content and flesh quality without the confounding effect of diet composition or nutritional status. After three generations of selection, fillets from F fish were consistently more colorful for both raw and cooked fillet, had a tendency to be less firm for raw fillet, but no difference was measured for cooked fillet texture evaluated by both instrumental and sensory methods (Lefevre et al., 2015). Those differences in quality parameters were associated with slightly larger white muscle fibers in F fish, especially in triploid fish (Lefevre et al., 2015).

As genetic selection is an effective way of controlling muscle fat content, we can assume that the difference in muscle fat content between L and F lines will increase with the number of generations of divergent selection. However, are there limit values for trout Fat-meter^®^ values, a minimum floor value for the L line, and a maximum ceiling value for the F line? Will the divergence in quality traits be of the same order as for fillet adiposity? These questions remained to be answered.

The objective of the present study was to compare some quality parameters and muscle features in fat muscle content divergently selected rainbow trout after seven generations of selection, in raw, cooked and smoked fillets.

## 2. Material and Methods

### 2.1. Ethical statement

PEIMA INRAE Fish Farming Systems Experimental Facility are authorized for animal experimentation under French regulation D29-277-02. The experiments were carried out from June 2013 to May 2014 following the Guidelines of the National Legislation on Animal Care of the French Ministry of Research (Decree N°2001-464, May 29, 2001). For these experiments, project authorization was not required. Experiments were conducted under the official license of Mrs Edwige QUILLET.

### 2.2. Fish and rearing conditions

Selected lines of rainbow trout, *Oncorhynchus mykiss*, were produced and reared in the INRAE’s experimental facilities (PEIMA, INRAE, Fish Farming systems Experimental Facility, DOI: 10.15454/1.5572329612068406E12, Sizun, France). Seven generations of divergent selection for high (F) or low (L) muscle fat content were performed using a non-destructive method (Distell Fish Fat-meter^®^) in live fish (Quillet et al., 2007). A correction was applied to Fat-meter values to take account of the positive correlation observed between absolute body weight and muscle lipid content (except in generations G3 and G4). The correction based on log-transformed weight was designed as the most appropriate (Quillet et al., 2007) until G7, where fish were selected on the basis of the deviation to (Fat/log(weight) regression. The mean pressure of selection per generation was 12.7% and 11.6% for F and L line, respectively.

The fish used in this study were reared in three replicated tanks per line until one year old, age of slaughter and stage corresponding to the size at which selection was performed. Fish were first in spring water at a steady temperature (≈ 11.5°C) for 1.5 months. Then, they were reared in circular 2 m diameter tanks containing 1.8 m^3^ water from the “Drennec” Lake (Sizun, France). The water temperature fluctuated seasonally from 7°C to 19°C. The water flow was adjusted to 2 tanks renewal per hour to allow an oxygen concentration above 6 mg/l and enable the disposal of fish waste. Routinely, 10% of the fish from each tank were weighed every 3 weeks. The fish were fed using successively continuous distribution system (1.5 month = spring water period), then 10 times (during 2.5 months) and 5 times a day (during 5 months) using automatic feeders, and finally with self-feeders (during the last 2 months). Diets (B Mega for organic trout production) were manufactured by an aquafeed producer (Le Gouessant, France) and were composed of 40% protein and from 24 to 28% lipids depending on pellet size. The ration level was calculated and adjusted each week. The amount of food distributed was increased by at least 10% compared to the usual ration tables to ensure that the fish were fed to satiation.

The fish were fasted for 48 h before slaughter and then harvested as fast as possible with a handling net, anesthetized with iso-eugenol (0.025ml/l) in a separate tank and bled by gill arch section. All the measurements at slaughter were performed immediately after death, within less than 1 h using 20 fish per replicate (60 fish per line). Ten fish per replicate (30 per line) were then measured for raw quality parameters at 48 h *post-mortem* and cooked quality parameters at 96 h *post-mortem*. Fillets from the other ten fish per replicate (30 per line) were then salted and smoked at 48h *post-mortem* and measured for smoked quality parameters at 7 days *post-mortem*.

### 2.3. Measurements at slaughter

The fish traits measurements were indexed according to the ontology ATOL (Animal Trait Ontology for Livestock, http://www.atol-ontology.com/index.php/en/les-ontologies-en/visualisation-en, (Golik et al., 2012).

The muscle lipid content (ATOL:0001663) assessment was conducted on whole fish using the Torry Fish Fat Meter^®^ (Distell Industries Ltd, Scotland). The fish were wiped with paper tissue to remove excess water and mucus. The instrument was firmly applied on the dorsal musculature, parallel to the lateral line, between the head and the dorsal fin of both sides of the fish (Douirin et al., 1998). The fat value was the mean of these two measurements.

Individual body weight (BW, ATOL:0000351) and standard length (L, ATOL:0001659) were measured. The condition factor (ATOL:0001653) was calculated as K=BW/L^3^. Body maximal width (Wi, in front of the dorsal fin) was measured and the ratio Wi/L was calculated.

The fish were eviscerated and filleted, sex was registered, and carcass, viscera, liver, gonads, head, and fillet were weighed. Carcass yield (= Carcass weight / BW), Hepato-Somatic Index (HSI = Liver weight / BW), and Gonado-Somatic Index (GSI = Gonad weight / BW) were calculated.

The fillet color (ATOL:0001017) was assessed using a portable Minolta Chromameter CR-400 (Minolta, France) equipped with light source C and a 2° observer angle, calibrated to a white standard. For each fillet, two measurements were taken on the interior part of the fillet, one anterior to the dorsal fin and the other anterior to the anal fin (**Figure 1**). The mean of these two measurements values was considered. Data were expressed using the L*, a*, b* system, representing lightness, redness, and yellowness, respectively, as recommended by CIE (CIE, 1976).

**Figure 1:**
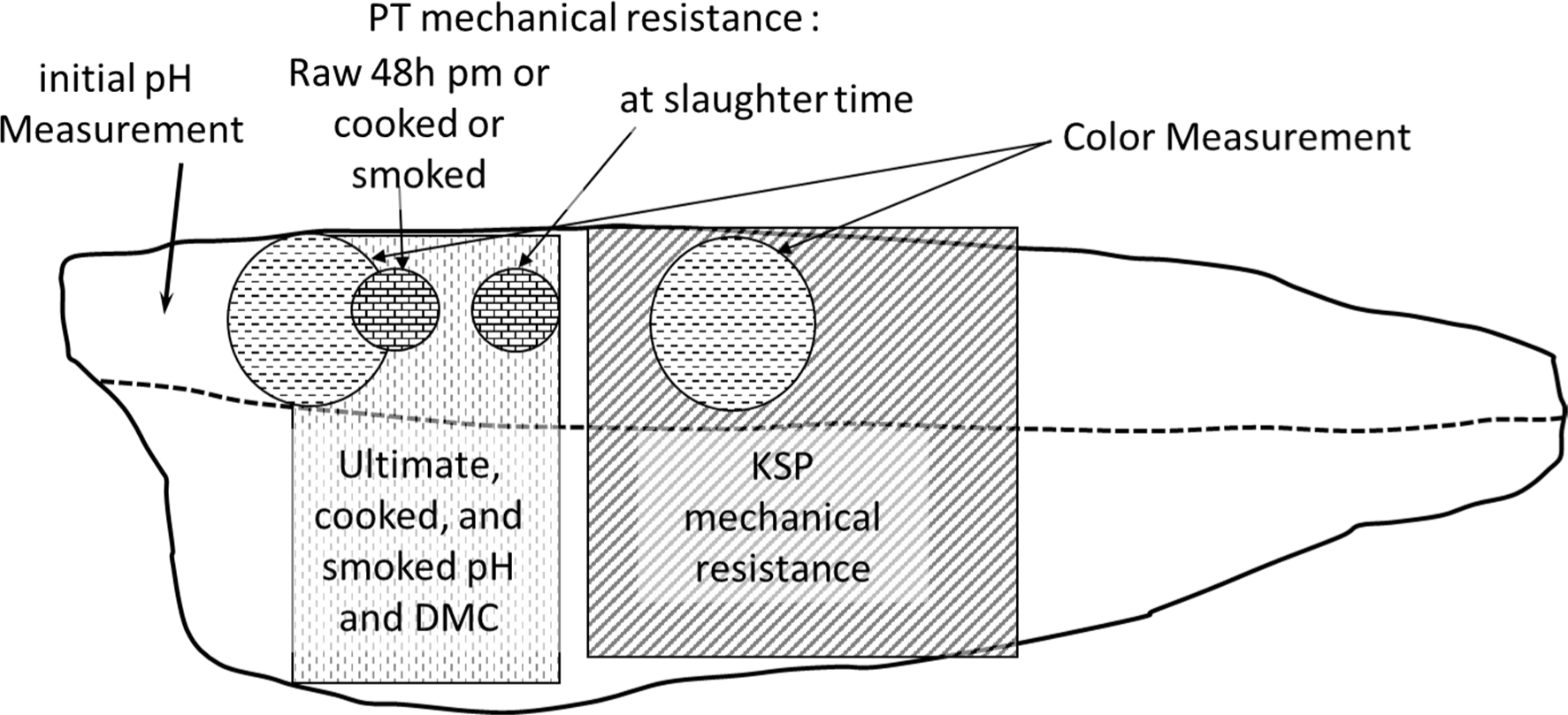
Schematic representation of the different measurements carried out on raw (at slaughter or at 48h *post-mortem*), cooked or smoked fillets; for details see text. PT = Penetrometry Test, KSP = Kramer Shear Press, DMC = Dry Matter Content.

The muscle initial pH (ATOL:0001684) was measured in the front part of the fillet within 30 min and 1 hour *post-mortem* using a Metrohm 826 mobile pH meter (Metrohm SA, Switzerland) equipped with a Metrohm spearhead electrode (Metrohm SA, Switzerland) and a Pt 1000 temperature sensor (Metrohm SA, Switzerland).

Right fillet mechanical resistance was evaluated with a penetrometry test in the dorsal part of the unskinned fillet, beneath the dorsal fin, just above the horizontal myoseptum (**Figure 1**). This test was done with a cylinder plunger (Ø 15 mm) mounted on a portable device (Andilog Textor B10, Andilog technologies, France), at a speed of 1 mm/sec until 2 mm from the support. The maximum force (N) and the total work of the test were registered.

Fillets were finally vacuum-packed within 1 hour after fish death and either stored on ice until further analysis at 2 days *post-mortem* (dpm, half of the fish) or stored in a cold-room (2°C) until salting and smoking at 2 dpm (the other half of the fish).

Muscle lipid content was measured on individual samples from 3 additional fish per tank (9 per line) as previously described (Kamalam et al., 2012).

### 2.4. Muscle histological analysis

Deep white muscle samples were taken within one hour after fish death, just beneath the dorsal fin. The samples were fixed in Carnoy fixative (absolute ethanol, chloroform, acetic acid, 6:3:1) for 48 h at 4°C, dehydrated in alcohol and alcohol/butanol and embedded in paraffin. Sample sections (10 μm) were then cut and stained with Sirius Red and Fast Green 0.1% in saturated picric acid (Lopez-De Leon and Rojkind, 1985). Five microscopic fields, presenting fibers with transversal sections, were digitized for each histological section. The areas of the transversal section of individual white muscle fibers (300-500 fibers per fish) were measured using Visilog 6.1 for Windows. Histological treatments, including paraffin embedding, cause muscle fiber shrinkage. Therefore, the individual muscle fiber area was multiplied by a shrinkage correction (SC) factor calculated as follows: SC = (total image area - connective tissue area) / (fiber total area)). The muscle fiber diameters (D) (ATOL:0000458) were then calculated using the formula D = 2√(area/π), under the assumption that the individual fiber cross-sections were circular.

### 2.5. Fillet Smoking

After two days of vacuum-packed storage in a cold room at 2°C, each individually marked fillet was weighed and “Sel de Guérande” salt (86% sodium chloride) was sprinkled on the flesh side of the unskinned fillets to cover the surface. Salting was carried out at 4°C, for 40 min. Then the fillets were rapidly rinsed, rested on grids at 4°C for approximately 1 hour. Weight of each fillet was recorded just before smoking. Fillet were then cold-smoked for 1 h at a mean temperature of 25◦C (24.5-25.4°C) with green beech wood in an air-conditioned and horizontally-ventilated smoking cabinet equipped with a GF 200 automatic smoke generator (ARCOS^®^ 01190 Gorrevod, France). At the end of the smoking, fillets were rested in the smoking chamber in cold air-conditioned temperature of 5°C. Smoked fillet were finally weighed, vacuum-packed, and stored at 4°C in a cold room, before refrigerated-transporting in expanded polystyrene (EPS) boxes with ice to the laboratory for further measurement. At the laboratory, smoked fillets were stored in a cold room at 4°C.

Salting yield (weight after salting / weight before salting) was 96.2 ± 0.5 % for fillets from L line vs 96.7 ± 0.7 % for fillets from F line (p<0.01). Salting + smoking yield (weight after salting+smoking / weight before salting+smoking) was 92.8 ± 1.0 % for fillets from L line vs 93.3 ± 0.9 % for fillets from F line (p<0.05).

### 2.6. Physical measurements of quality parameters of raw (2 dpm), cooked (4 dpm) and smoked (7 dpm) fillet

All the physical measurements were done at room temperature. At 2 days *post-mortem* (dpm), physical measurements of the quality parameters were performed on one raw fillet, whereas the second fillet was cooked. The fillet mean weight was approximately 75 g. Skinless fillets were cooked for 1 to 2 min, depending on their weight, in a domestic microwave oven (Samsung M192DN) at 450 W in a covered bowl to reach a core temperature of 65°C-70°C. Fillets were then cooled to room temperature, weighed, packed in plastic bags and stored at 4°C in a cold room until further analysis. Cooking yield (weight after cooking / weight before cooking) was 84.3 ± 1.4 % for fillets from L line vs 83.8 ± 1.2 % for fillets from F line (p>0.05). Physical measurements of the quality parameters for the cooked fillets were assessed at 96 h *post-mortem*.

The fillet color was measured as previously described for the measurement at slaughter (Section 2.2).

The ultimate pH (ATOL:0001684) was measured with 5 g muscle, sampled in the anterior part of the fillet (**Figure 1**) and homogenized in four volumes of distilled water.

The dry matter content (ATOL:0000101) was determined by drying approximately 5 g of minced fillet (anterior part, **Figure 1**) for 40 h in an oven at 105°C.

The mechanical strength (ATOL:0001649) of the middle part of the skinned fillet (64 mm length beneath the dorsal fin) was analyzed using a Kramer shear cell mounted on a static load cell of 2 kN (INSTRON 5544, INSTRON Ltd., England). The fillet sample was completely sheared with shear blades perpendicular to the main axis of the fish (perpendicular to muscle fibers) at a speed of 1 mm/sec. The maximum force (N) and the work until the maximum force were calculated. These parameters were divided by the weight of the sample (Szczesniak et al., 1970).

A penetrometry test, similar to those described under the slaughter methods (Section 2.2), was done on skinned fillet just in front of the slaughter measurement (Figure 1).

### 2.7. Statistical analysis

All of the results are expressed as the mean ± standard deviation. For all the parameters, a two-ways ANOVA was used to test the effects of line (L vs F) and sex (female vs male). As no effect of sex was measured for almost all parameters, the results are presented as a result of an ANOVA comparing L and F lines. When the effect of sex was significant, the result of the two ways ANOVA are reported (**Table 1**). The significant level was set at p<0.05. The significant differences between the mean values were determined using the Newman-Keuls test. The Pearson correlation coefficient was calculated to analyze the significance of the linear relationships between variables. All of the analyses were performed using Statistica for Windows (version 5.1) software. The number of fishes measured, from 9 to 60, for each parameter is specified in the tables.

**Table 1:**
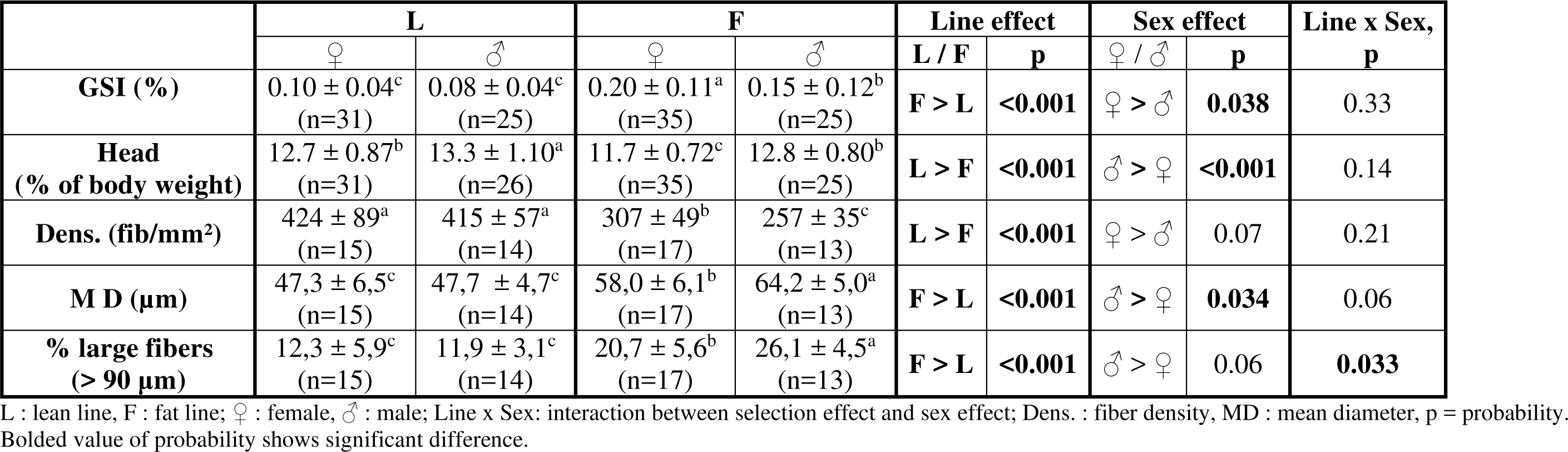
Effects of divergent selection for muscle lipid content and sex on some fish and white muscle fibers characteristics. Mean ± standard deviation.

|  | **L** |  | **F** |  | **Line effect** |  | **Sex effect** |  | **Line x Sex,**<br>**p** |
| --- | --- | --- | --- | --- | --- | --- | --- | --- | --- |
|  | ♀ | ♂ | ♀ | ♂ | **L / F** | **p** | ♀ / ♂ | **p** |  |
| **GSI (%)** | 0.10 ± 0.04<sup>c</sup><br>(n=31) | 0.08 ± 0.04<sup>c</sup><br>(n=25) | 0.20 ± 0.11<sup>a</sup><br>(n=35) | 0.15 ± 0.12<sup>b</sup><br>(n=25) | **F > L** | **<0.001** | ♀ > ♂ | **0.038** | 0.33 |
| **Head**<br>**(% of body weight)** | 12.7 ± 0.87<sup>b</sup><br>(n=31) | 13.3 ± 1.10<sup>a</sup><br>(n=26) | 11.7 ± 0.72<sup>c</sup><br>(n=35) | 12.8 ± 0.80<sup>b</sup><br>(n=25) | **L > F** | **<0.001** | ♂ > ♀ | **<0.001** | 0.14 |
| **Dens. (fib/mm<sup>2</sup>)** | 424 ± 89<sup>a</sup><br>(n=15) | 415 ± 57<sup>a</sup><br>(n=14) | 307 ± 49<sup>b</sup><br>(n=17) | 257 ± 35<sup>c</sup><br>(n=13) | **L > F** | **<0.001** | ♀ > ♂ | 0.07 | 0.21 |
| **M D (μm)** | 47,3 ± 6,5<sup>c</sup><br>(n=15) | 47,7 ± 4,7<sup>c</sup><br>(n=14) | 58,0 ± 6,1<sup>b</sup><br>(n=17) | 64,2 ± 5,0<sup>a</sup><br>(n=13) | **F > L** | **<0.001** | ♂ > ♀ | **0.034** | 0.06 |
| **% large fibers**<br>**(> 90 μm)** | 12,3 ± 5,9<sup>c</sup><br>(n=15) | 11,9 ± 3,1<sup>c</sup><br>(n=14) | 20,7 ± 5,6<sup>b</sup><br>(n=17) | 26,1 ± 4,5<sup>a</sup><br>(n=13) | **F > L** | **<0.001** | ♂ > ♀ | 0.06 | **0.033** |

L : lean line, F : fat line; ♀ : female, ♂ : male; Line x Sex: interaction between selection effect and sex effect; Dens. : fiber density, MD : mean diameter, p = probability. Bolded value of probability shows significant difference.

A multiple linear regression using the RegBest function of the FactoMineR of R software package was performed using flesh color L*a*b* measurement, muscle initial and ultimate pH, raw dry matter content, fat meter value and muscle fiber size (density, mean diameter, percentage of small and large fiber) as independent variables and the raw fillet specific resistance as dependent variable.

A Principal Component Analysis (PCA) was also applied, using Factoshiny package of R, with adiposity, raw fillet quality, and muscle cellularity parameters.

## 3. Results

### 3.1. Fish characteristics

Main fish characteristics are presented in **Table 2**. No difference of body weight was measured between F and L lines. However, fish from L line were longer (+ 5%), and as a consequence had a lower Fulton condition factor values (1.47±0.16 *vs* 1.62±0.13 p<0.001 for L and F lines, respectively). Consistently, fish from the F line were wider, both in measured value and even more when the width was expressed as a percent of fish length. Fish from the F line also had a lower relative head weight compared to fish from the L line.

**Table 2:** Effects of divergent selection for muscle lipid content on fish characteristics.

|  | L |  |  | F |  |  | L / F | Anova<br>p |
| --- | --- | --- | --- | --- | --- | --- | --- | --- |
|  | mean | sd | n | mean | sd | n |  |  |
| Fish morphology |  |  |  |  |  |  |  |  |
| Weight (g) | 382 | 73 | 60 | 362 | 71 | 60 |  | 0.12 |
| Standard Length (mm) | 296 | 14 | 60 | 281 | 20 | 60 | L > F | <0.001 |
| K | 1.47 | 0.16 | 60 | 1.62 | 0.13 | 60 | F > L | <0.001 |
| Fish maximal width (mm) | 35 | 3 | 60 | 37 | 3 | 60 | F > L | 0.003 |
| Width/Length (%) | 11.9 | 0.72 | 60 | 13.2 | 0.78 | 60 | F > L | <0.001 |
| Yields |  |  |  |  |  |  |  |  |
| Carcass yield (%) | 85.2 | 2.1 | 60 | 84.9 | 1.4 | 60 |  | 0.31 |
| HSI (%) | 1.17 | 0.21 | 60 | 1.44 | 0.26 | 59 | F > L | <0.001 |
| GSI (%) | 0.09 | 0.04 | 57 | 0.17 | 0.12 | 60 | F > L<sup>§</sup> | <0.001 |
| Head (% of body weight) | 13.1 | 1.05 | 60 | 12.1 | 0.92 | 60 | L > F<sup>§</sup> | <0.001 |
| Raw fillet (% of body weight) | 61.3 | 2.25 | 60 | 62.5 | 2.61 | 60 | F > L | 0.008 |
| Dressed/Skinned Fillet (% of body weight)<sup>§§</sup> | 43.3 | 2.6 | 30 | 45.6 | 4.0 | 30 | F > L | 0.012 |
| Adiposity |  |  |  |  |  |  |  |  |
| Fat-meter® Measurement (%) | 5.06 | 0.81 | 60 | 20.9 | 4.8 | 60 | F > L | <0.001 |

L : lean line, F : fat line; sd = standard deviation; n = number of fish measured; p = probability. Bolded value of probability shows significant difference; HIS = Hepato-Somatic Index = Liver Weight / Body Weight; GSI = Gonado-Somatic Index = Gonad Weight / Body Weight; § : Sex effect $p < 0.05$ , see Table 2 for the result of two-ways ANOVA; §§ : Calculated after weighing dressed/skinned filler at 48h *post-mortem*.

No difference of carcass yield was measured between the two lines but fish from F line had higher hepato-somatic index, gonado-somatic index, and fillet index.

As expected, fish from the F line exhibited much higher Fat-meter^®^ values than fish from the L line (20.9±4.8 *vs* 5.06±0.81 p<0.001 for F and L lines, respectively).

A difference between males and females (**Table 1**) was measured for gonado-somatic index, males giving lower value than females, and the relative weight of fish head, males giving higher value than female, without significant interaction between line and sex.

### 3.2. Characteristics of raw, cooked and smoked fillets

As expected, muscle lipid content was higher in F fish compared to L fish (13.5±2.2 vs 5.0±1.0% of lipid for F and L fish, respectively). Consistently dry matter content of fillet was much higher for F fish than for L fish both for raw (32.3±2.1 vs 25.5±1.3% of fillet dry matter content for F and L fish, respectively), cooked, and smoked fillets (**Table 3**).

**Table 3:** Effects of divergent selection for muscle lipid content on pH, color parameters and adiposity of raw, cooked and smoked fillets.

|  | L |  |  | F |  |  | L / F | Anova<br>p |
| --- | --- | --- | --- | --- | --- | --- | --- | --- |
|  | mean | sd | n | mean | sd | n |  |  |
| Raw fillet |  |  |  |  |  |  |  |  |
| Lipid content (%) | 5.02 | 1.02 | 9 | 13.51 | 2.24 | 9 | F > L | <0.001 |
| DMC (%) | 25.5 | 1.3 | 30 | 32.3 | 2.1 | 30 | F > L | <0.001 |
| pH slaughter | 7.09 | 0.14 | 60 | 7.12 | 0.10 | 60 |  | 0.31 |
| pH 48h pm | 6.37 | 0.04 | 30 | 6.35 | 0.05 | 30 |  | 0.10 |
| L* slaughter | 46.5 | 1.73 | 60 | 48.8 | 1.85 | 60 | F > L | <0.001 |
| L* 48h pm | 44.5 | 1.9 | 30 | 47.4 | 1.5 | 30 | F > L | <0.001 |
| a* slaughter | 0.55 | 0.48 | 60 | 0.81 | 0.35 | 60 | F > L | <0.001 |
| a* 48h pm | 0.26 | 0.41 | 30 | 0.45 | 0.37 | 30 |  | 0.06 |
| b* slaughter | 1.52 | 1.12 | 60 | 3.12 | 0.93 | 60 | F > L | <0.001 |
| b* 48h pm | -0.40 | 0.75 | 30 | 1.69 | 1.09 | 30 | F > L | <0.001 |
| Cooked fillet |  |  |  |  |  |  |  |  |
| DMC cooked (%) | 29.8 | 1.2 | 30 | 35.9 | 1.5 | 30 | F > L | <0.001 |
| pH cooked | 6.62 | 0.04 | 30 | 6.61 | 0.04 | 30 |  | 0.71 |
| L* cooked | 76.3 | 1.7 | 30 | 76.0 | 2.1 | 30 |  | 0.57 |
| a* cooked | 0.21 | 0.61 | 30 | 0.33 | 0.49 | 30 |  | 0.39 |
| b* cooked | 13.9 | 1.3 | 30 | 14.8 | 1.1 | 30 | F > L | 0.005 |
| Smoked fillet |  |  |  |  |  |  |  |  |
| DMC smoked (%) | 28.4 | 1.2 | 30 | 34.6 | 2.2 | 30 | F > L | <0.001 |
| pH smoked | 6.18 | 0.04 | 30 | 6.18 | 0.04 | 30 |  | 0.86 |
| L* smoked | 46.3 | 1.3 | 30 | 46.4 | 1.1 | 30 |  | 0.82 |
| a* smoked | -0.76 | 0.25 | 30 | -0.27 | 0.38 | 30 | F > L | <0.001 |
| b* smoked | 0.44 | 1.26 | 30 | 2.04 | 1.20 | 30 | F > L | <0.001 |

L : lean line, F : fat line; sd = standard deviation; n = number of fish measured; p = probability. Bolded value of probability shows significant difference; L\*, a\*, b\* = color parameters of the fillet, L\* : lightness, a\* : redness, b\* : yellowness; DMC : Dry Matter Content.

No difference of muscle pH was measured between lines both at slaughter or 48 h pm for raw fillet, nor for cooked and smoked fillets (**Table 3**). Raw fillet lightness (L*) was higher for F fish that for L fish both at slaughter and at 48 h pm. However, this difference is no longer observed for cooked or smoked fillets. As fish were fed a non-pigmented diet, values of redness and yellowness were very low. Nevertheless, higher value of redness (a*), at slaughter time for raw fillet and for smoked fillets, and yellowness (b*), for both, raw, cooked and smoked fillets, were measured for F fish fillets than for L ones.

### 3.3. Mechanical resistance of raw, cooked and smoked fillets

Higher values of mechanical resistance were measured (**Table 4**) for raw fillet from L fish compared to F fish, both at slaughter time (< 2 h pm) and 48 h pm, with the two measurement tools (cylinder plunger and Kramer shear press). For example, specific resistance (Fmax / sample weight) measured in Kramer shear press test was 21% higher for raw fillet from the L line compared to fillet from the F line.

**Table 4:** Effects of divergent selection for muscle lipid content on mechanical resistance of raw, cooked and smoked fillets raw, cooked and smoked fillets.

| Method | Parameter : | L |  |  | F |  |  | L / F | Anova<br>p |
| --- | --- | --- | --- | --- | --- | --- | --- | --- | --- |
|  |  | mean | sd | n | mean | sd | n |  |  |
| Raw fillet at slaughter |  |  |  |  |  |  |  |  |  |
| PT | Fmax (N) | 37.7 | 4.93 | 60 | 33.4 | 4.31 | 60 | L > F | <0.001 |
|  | W/wi (AU) | 96.3 | 8.77 | 60 | 90.0 | 9.08 | 60 | L > F | <0.001 |
| Raw fillet at 48h *post-mortem* |  |  |  |  |  |  |  |  |  |
| PT | Fmax (N) | 10.5 | 1.8 | 30 | 9.1 | 1.2 | 30 | L > F | <0.001 |
|  | W/wi (AU) | 27.0 | 4.9 | 30 | 27.8 | 5.1 | 30 |  | 0.54 |
| KSP | Fmax (N) | 433 | 61 | 30 | 352 | 53 | 30 | L > F | <0.001 |
|  | Fmax/we (N/g) | 12.7 | 1.9 | 30 | 10.0 | 1.1 | 30 | L > F | <0.001 |
| Cooked Fillet |  |  |  |  |  |  |  |  |  |
| PT | Fmax (N) | 37.1 | 3.7 | 30 | 36.7 | 4.0 | 30 |  | 0.66 |
|  | W/wi (AU) | 67.2 | 7.9 | 30 | 63.6 | 7.7 | 30 |  | 0.080 |
| KSP | Fmax (N) | 1086 | 168 | 30 | 1097 | 162 | 30 |  | 0.78 |
|  | Fmax/we (N/g) | 36.4 | 5.4 | 30 | 34.5 | 3.8 | 30 |  | 0.12 |
| Smoked Fillet |  |  |  |  |  |  |  |  |  |
| PT | Fmax (N) | 20.7 | 2.1 | 30 | 18.3 | 3.2 | 30 | L > F | 0.001 |
|  | W/wi (AU) | 57.0 | 4.5 | 30 | 55.7 | 7.9 |  |  | 0.47 |
| KSP | Fmax (N) | 482 | 79 | 30 | 429 | 57 | 30 | L > F | 0.004 |
|  | Fmax/we (N/g) | 16.4 | 3.0 | 30 | 14.8 | 2.7 | 30 | L > F | 0.035 |

L : lean line, F : fat line; sd = standard deviation; n = number of fish measured; p = probability. Bolded value of probability shows significant difference; PT : Puncture Test; KSP : Kramer Shear Press, Fmax : maximal force, W : Work; .wi : sample width; we : sample weight; AU : Arbitrary Unit.

A difference between the two lines was also measured for smoked fillet (16.4±3.0 vs 14.8±2.7 N/g, p<0.05, for specific resistance of L and F lines, respectively), but no significant difference between fillets from fish of the two lines were measured for cooked fillet (p>0.05).

### 3.4 Muscle histological analysis

A large difference in white muscle fiber size was observed between the two lines (**Table 5**). Mean diameter of white muscle fiber of F fish was 22% higher than this of L fish. Consequently, white muscle fiber density was higher for L fish compared to F fish (419±73 vs 285±49 fibers/mm^2^, p<0.001, for L and F fish, respectively). More precisely, L fish had almost twice the amount of small (<20 µm) fibers (19±6% *vs* 10±4%, p<0.001, for L and F fish, respectively) and less large (>90 µm) fibers (12±5 vs 23±6%, p<0.001, for L and F fish, respectively) than F fish.

**Table 5:** Effects of divergent selection for muscle lipid content on white muscle fiber density (Dens.), mean diameter (MD), percentage of small fibers (diameter < 20 μm), and percentage of large fibers (diameter > 90 μm).

|  | **L** |  |  | **F** |  |  | **L / F** | **Anova** |
| --- | --- | --- | --- | --- | --- | --- | --- | --- |
|  | **mean** | **sd** | **n** | **mean** | **sd** | **n** |  | **p** |
| **Dens. (fib/mm<sup>2</sup>)<sup>§</sup>** | 419 | 73 | 30 | 285 | 49 | 30 | **L > F** | **<0.001** |
| **M D (µm)<sup>§</sup>** | 47.5 | 5.5 | 30 | 60.7 | 6.4 | 30 | **F > L** | **<0.001** |
| **% small fibers (< 20 µm)** | 19.3 | 5.5 | 30 | 9.7 | 3.8 | 30 | **L > F** | **<0.001** |
| **% large fibers<sup>§</sup> (> 90 µm)** | 12.1 | 4.6 | 30 | 23.0 | 5.7 | 30 | **F > L** | **<0.001** |

An effect of sex was observed for white muscle fiber density, mean diameter, and the percent of large fibers (**Table 2**). Males had a higher white muscle fiber mean diameter (p<0.05), and a tendency (p<0.10) to have a lower fiber density and a higher percent of large fibers. A significant interaction between line and sex effects was measured for the percent of large fibers: no difference between males and females was observed for fish of the L line whereas males from the F line had a higher proportion of large fibers that females (26±5 *vs* 21±6% for males and females of the F line, respectively).

### 3.5. Correlations, multiple linear regression, and PCA analyses

Pearson correlation coefficient was calculated for fillet color parameters (**Table S1**), mechanical resistance parameters (**Table S2**), and to compare smoked fillet characteristics to their raw counterpart (**Table S3**).

For raw fillet, lightness (L*) and yellowness (b*) were positively correlated to muscle adiposity (r=0.61 for L*r, r=0.69 for b*r, respectively, and dry matter content, p<0.001) and white muscle fiber size (r=0.65 for L*s, r=0.53 for b*s, respectively, and mean fiber diameter, p<0.001) (**Figure 2**). For cooked fillet, the same trend was observed for yellowness but no longer for lightness. Cooked fillet lightness was not correlated to that of raw fillet whereas cooked fillet redness and yellowness were positively correlated to those of raw fillet (**Table S1**).

**Figure 2:**
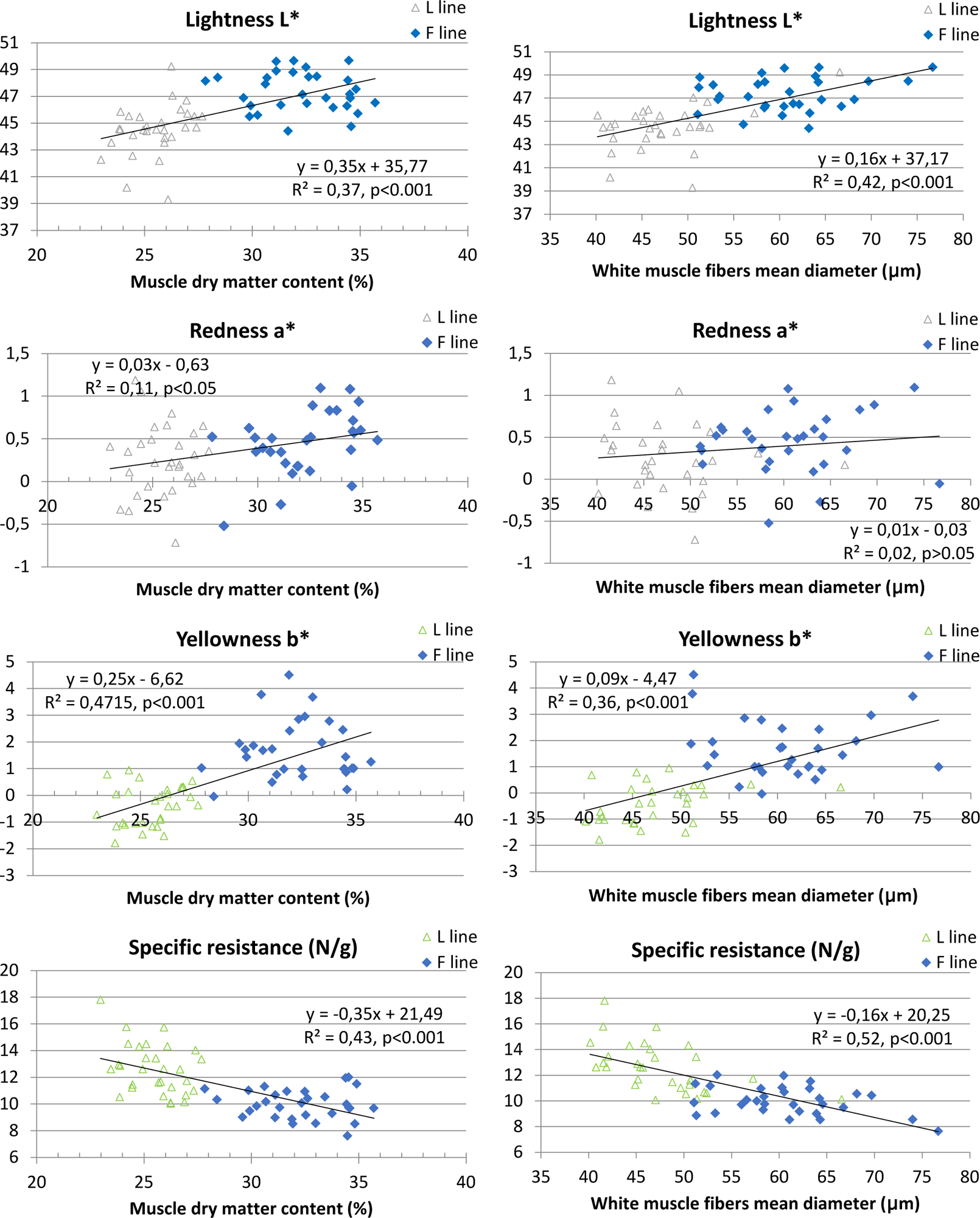
Pearson correlation between **raw fillet** instrumentally measured quality parameters and muscle dry matter content and mean fibers diameter, L line: green open triangle, F line: full blue diamond, n=60.

Mechanical resistance parameters or raw fillet measured at slaughter were correlated to those measured at 48 h pm (**Table S2**). However, mechanical resistance parameters for cooked fillet were not strongly correlated to those of raw fillet. Mechanical resistance parameters of raw fillet were strongly negatively correlated to muscle adiposity and to white muscle fiber size (r=-0.66 for Kramer Shear Press specific resistance and muscle dry matter content, and r=-0.72 for Kramer Shear Press specific resistance and white muscle fiber mean diameter, p<0.001) (**Figure 2**).

Most of quality parameters measured for smoked fillet were positively correlated to raw fillet ones (**Figure 3**). For example, maximum force measured in penetrometry test measured for smoked fillet at 7 days *post-mortem* was correlated to that measured for raw fillet at slaughter (r=0.69, p<0.001). However, we can notice that smoked fillet lightness was not significantly correlated to that of raw fillet (r=0.25, p>0.05), whereas redness and yellowness were.

**Figure 3:**
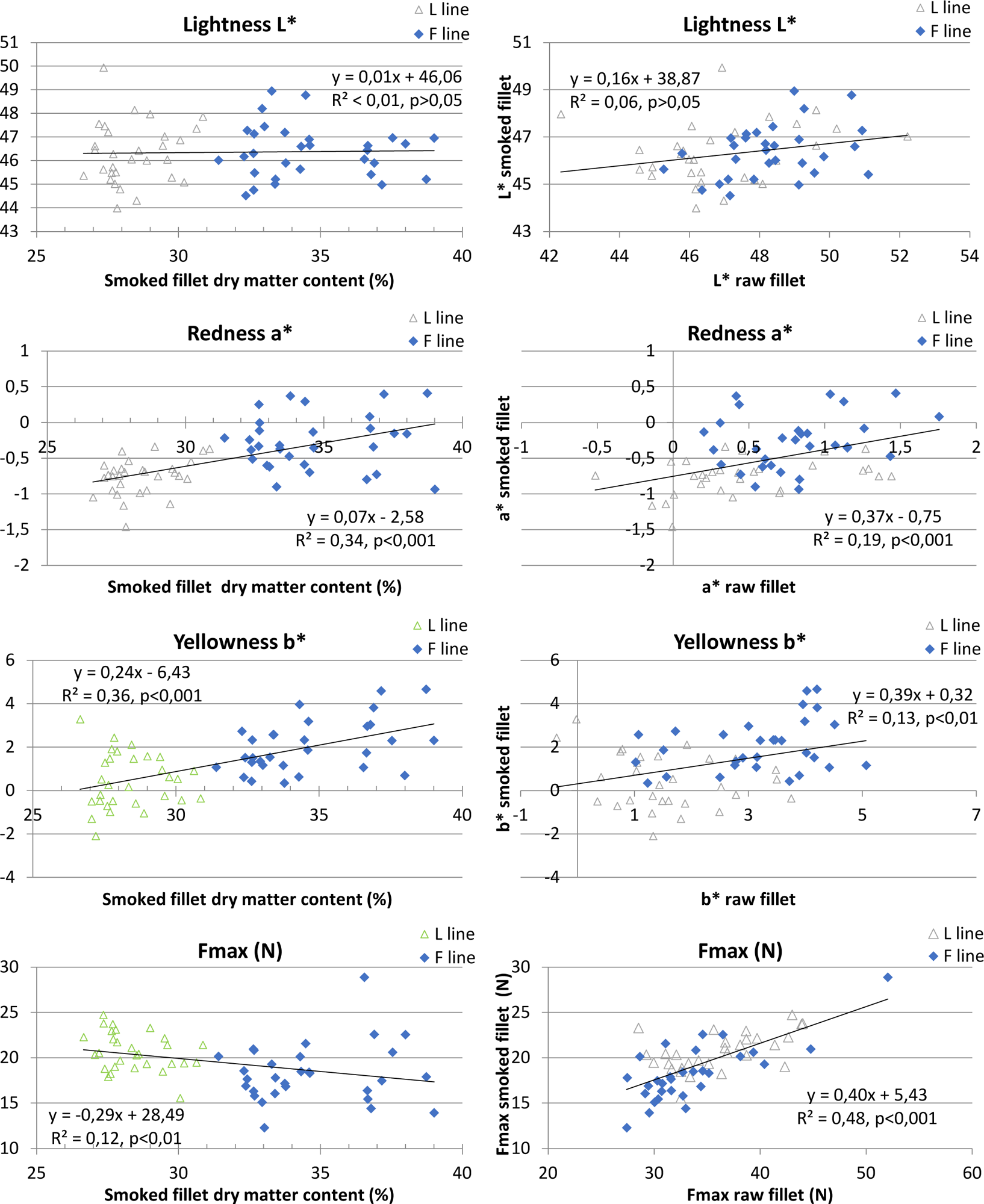
Pearson correlation between **smoked fillet** instrumentally measured quality parameters and smoked fillet dry matter content and between smoked and raw fillet quality parameters, L line: green open triangle, F line: full blue diamond, n=60.

Moreover, as for raw fillet, smoked fillet yellowness b*, but also redness a*, was correlated to fillet adiposity (r=0.59, p<0.001 for both a* and b* with fat-meter® value). However, smoked fillet mechanical resistance was less strongly negatively correlated to muscle adiposity than raw fillet one. Maximal value was r=-0.35 (p<0.01) for maximal force in penetrometry test and smoked fillet dry matter content).

The multiple linear regression shows that the best model can significantly predict specific mechanical resistance of the raw flesh (R²=0.68, p<0.001) with four variables (fiber density, percentage of large fiber, fillet yellowness (b*) at slaughter, and ultimate pH).

The PCA plot (**Figure 4**) shows that the two lines were separated on the first axis comprising more than 45% of the variability. This analysis also sums up the positive association of both adiposity and muscle fiber size with fillet color parameters, and the negative one with raw fillet mechanical resistance.

**Figure 4:**
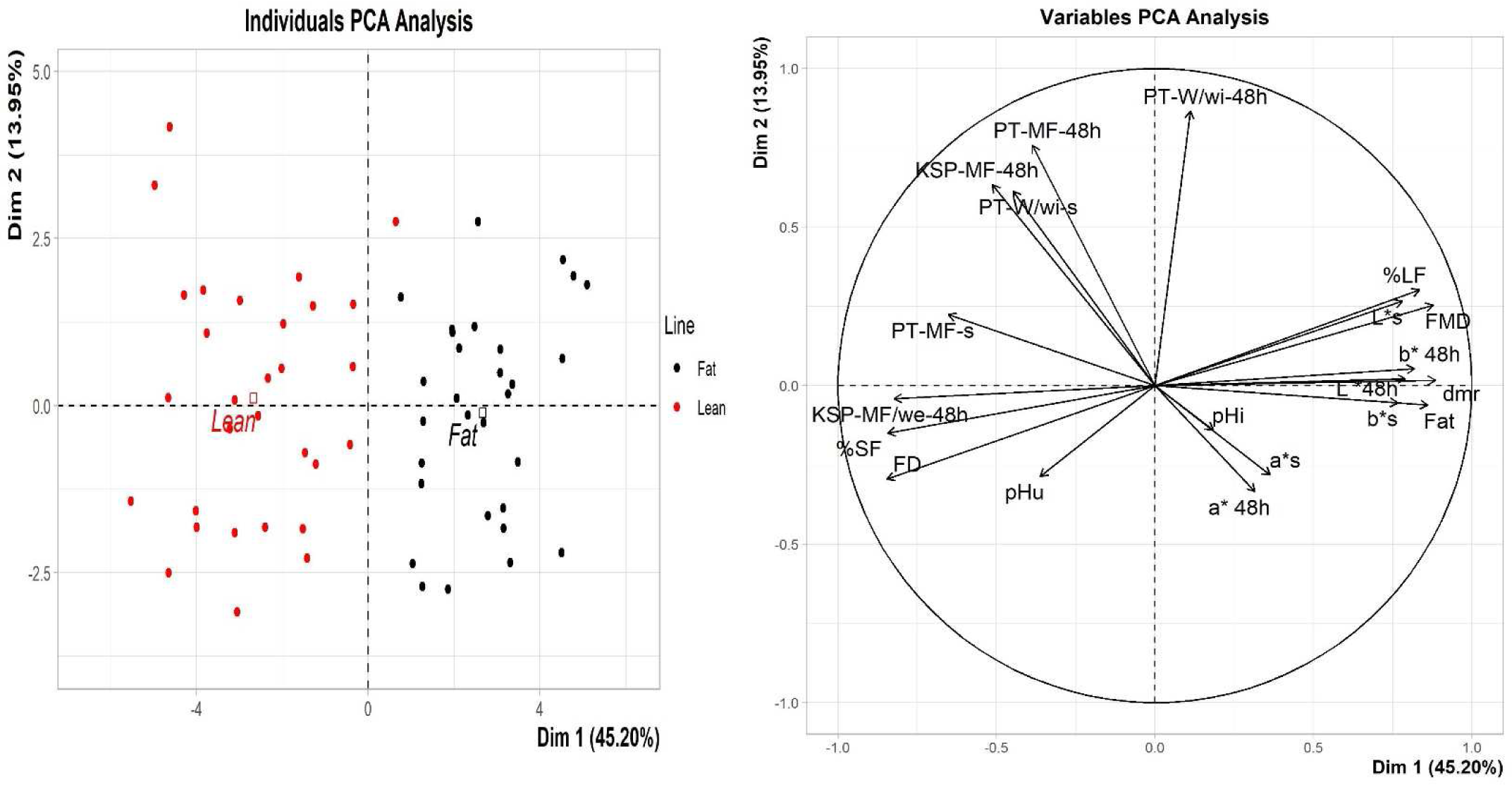
Principal Component Analysis (PCA) of adiposity, raw fillet quality, and muscle cellularity parameters, n=60. Fat: Fat-meter value; dmr: raw muscle dry matter content; pHi: initial pH; pHu: ultimate pH; L*s, a*s, b*s: lightness, redness, yellowness at slaughter; L* 48h, a* 48h, b* 48h: lightness, redness, yellowness for raw flesh at 48h *post-mortem*; PT: Puncture Test; KSP: Kramer Shear Press, MF: maximal force, wi: sample width; we: sample weight; ‘s’: measurement at slaughter; ‘18h’: raw fillet measured at 48h *post-mortem*; FD: Fiber density; FMD: Fiber Mean Diameter; % SF: % of Small Fibers; %LF: % of Large Fibers.

## 4. Discussion

The objective of the present work was to evaluate quality parameters in raw, cooked and smoked rainbow trout after seven generations of divergent selection on adiposity measured by Distell Fish Fat-meter^®^ (Quillet et al., 2005).

### 4.1. From 3^rd^ to 7^th^ generation of selection: divergence of adiposity leads to divergence in textural parameters and even more in muscle cellularity

The effects of divergent selection for muscle fat content were already evaluated for pan-size diploid and triploid fish from the third generation of selection for raw and cooked fillet (Lefevre et al., 2015) (see **Table 6** for comparison between generations). The present work reinforces the value of this genetic model of differential muscle adiposity to shed light on the consequences of adiposity on product qualities. First, a great increase of mean fat-meter value was observed between the third and the seventh generation for F line whereas only a small decrease was measured for L line. Body weight was not different between the two lines but fish from F line confirmed their more “compact” morphology demonstrated by lower body length, higher condition factor value and higher body width (Lefevre et al., 2015). Carcass yield was similar between the two lines, which differ from previous work reporting a higher viscero-somatic index for L line (Lutfi et al., 2018). Fish from F line also had higher hepato-somatic (HSI) and gonado-somatic (GSI) indexes. Even if not observed in younger stage, higher HSI in F line can be related to its higher lipogenic activity, fatty acid bioconversion and higher liver glycogen content (Kamalam et al., 2012). These discrepancies between different studies show that some phenotypic differences between lines are well established and robust, while others are less consistently observed, like higher body weight, higher perivisceral fat deposits leading to lower carcass yield, or lower HSI, for the L line. It seems therefore that the expression of these phenotypes is more dependent on the rearing environment, including the ration level, the type of the feed, or the thermal profile which can vary from one year to another. Phenotypic differences that are systematically observed include the morphology (higher value of condition factor or smaller head for F line for example) or the higher GSI in the F line. This higher GSI for F line, even before starting sexual maturation, was already observed in previous generations, as was precocious puberty (Weil et al., 2008). On the other hand, the higher relative muscle mass of F line, revealed by higher fillet yields, can be related to the difference of fish morphology, as negative genetic correlation was observed between fillet yield and bony tissue development as head proportion (Haffray et al., 2012). By the way, the higher body length and the higher head development of the L line highly suggest a greater skeletal development in this line. Moreover, the larger muscle fibers observed for F line suggest a higher muscle hypertrophy, i.e. growth by increasing muscle fibers size, in that line which could contribute to more muscle mass.

**Table 6:** Comparison of the effects of 3^rd^ and 7^th^ generation of divergent selection for muscle fat content on fish characteristics, fillet quality parameters and white muscle histological analysis. Mean ± standard deviation, n ≥ 75 for G3 fish traits and slaughter quality parameters, n = 60 for G7 fish traits and slaughter quality parameters, n=15 for G3 raw 48h *post-mortem* fillet, cooked fillet quality parameters and histological data, n=30 for G7 raw 48h *post-mortem* fillet, cooked fillet quality parameters and histological data

|  | L |  | F |  | G3 effect | G3 effect | G7 effect | G7 vs G3 | G7 vs G3 | F vs L | F vs L |
| --- | --- | --- | --- | --- | --- | --- | --- | --- | --- | --- | --- |
| | G3 | G7 | G3 | G7 | 2n+3n<sup>\$</sup> | 2n | | L | F | G3 | G7 |
| **Fish traits** |  |  |  |  |  |  |  |  |  |  |  |
| **BW (g)** | 302±66 | 382±73 | 290±55 | 362±71 | NS | NS | NS | +26.6% | +24.8% | -4.10% | -5.40% |
| **K** | 1.76±0.15 | 1.47±0.16 | 1.82±0.19 | 1.62±0.13 | ** | ** | *** | -16.6% | -11.3% | +3.72% | +10.3% |
| **Fat-meter<sup>®</sup> (%)** | 5.97±0.90 | 5.06±0.81 | 11.5±1.8 | 20.9±4.8 | *** | *** | *** | -15.3% | +82.8% | +91.9% | +314% |
| **Raw fillet quality parameters** |  |  |  |  |  |  |  |  |  |  |  |
| **L\* slaughter<sup>(1)</sup>** | 43.3±2.0 | 46.6±1.7 | 44.2±1.8 | 48.8±1.8 | *** | ** | *** | ND | ND | +7.58% | +10.2% |
| **a\* slaughter<sup>(1)</sup>** | 9.2±1.5 | 0.55±0.48 | 9.9±1.2 | 0.81±0.35 | *** | *** | *** | ND | ND | -94.0% | -91.2% |
| **b\* slaughter<sup>(1)</sup>** | 14.4±1.8 | 1.52±1.12 | 15.1±1.7 | 3.12±0.93 | *** | ** | *** | ND | ND | -89.4% | -79.3% |
| **DMC (%)** | 27.0±1.1 | 25.5±1.3 | 29.6±1.5 | 32.3±2.1 | *** | *** | *** | -5.72% | +9.08% | +9.47% | +26.7% |
| **KSP-Fmax (N)** | 387±51 | 433±61 | 351±38 | 352±53 | *** | * | *** | +11.9% | +0.04% | -9.14% | -18.9% |
| **KSP-Fmax/we (N/g)** | 13.2±2.1 | 12.7±1.9 | 12.2±1.9 | 10.0±1.1 | * | NS | *** | -4.08% | -18.3% | -7.28% | -21.0% |
| **Cooked fillet quality parameters** |  |  |  |  |  |  |  |  |  |  |  |
| **DMC (%)** | 28.8±1.5 | 29.8±1.2 | 30.0±1.9 | 35.9±1.5 | *** | p=0.054 | *** | +3.63% | +19.7% | +4.37% | +20.6% |
| **KSP-Fmax (N)** | 572±95 | 1086±168 | 630±148 | 1097±162 | NS | NS | NS | +89.9% | +74.1% | +10.3% | +1.08% |
| **KSP-Fmax/we (N/g)** | 25.8±4.4 | 36.4±5.4 | 28.3±6.8 | 34.5±3.8 | NS | NS | NS | +41.0% | +21.7% | +9.89% | -5.16% |
| **White muscle histological data** |  |  |  |  |  |  |  |  |  |  |  |
| **Dens. (fib/mm<sup>2</sup>)** | 470±79 | 419±73 | 442±70 | 285±49 | * | NS | *** | -10.9% | -35.4% | -6.07% | -31.9% |
| **M D (μm)** | 45.4±3.9 | 47.5±5.5 | 46.3±3.9 | 60.7±6.4 | * | NS | *** | +4.6% | +31.0% | +2.12% | +27.8% |
| **% small fibers (< 20μm)** | 22.6±5.4 | 19.3±5.5 | 21.8±5.1 | 9.7±3.8 | * | NS | *** | -14.5% | -55.8% | -3.13% | -49.9% |
| **% large fibers (> 100μm)** | 7.6±5.0 | 12.1±4.6 | 9.3±4.8 | 23.0±5.7 | p=0.08 | NS | *** | +58.6% | +147.7% | +21.7% | +90.0% |

L : lean line, F : fat line. G3 : 3<sup>rd</sup> generation of selection, G7 : 7<sup>th</sup> generation of selection; 2n : diploid fish, 3n : triploid fish; BW: body weight; K: condition factor; Fat: Fat-meter value; L\* : lightness, a\* : redness, b\* : yellowness, (1) : Fish evaluated for the 3<sup>rd</sup> generation of selection had received a fed pigmented diet whereas unpigmented diet was used for the rearing of the 7<sup>th</sup> generation. DMC : Dry Matter Content; KSP : Kramer Shear Press, Fmax : maximal force, we : sample weight; Dens. : white muscle fiber density, MD : mean diameter; NS : not significant ( $p>0.05$ ), \* : $p<0.05$ , \*\* : $p<0.01$ , \*\*\* : $p<0.001$ . \$ : Lefèvre et al., 2015. The ratios were calculated as follows : F vs L = (Fvalue-Lvalue)/Lvalue, G7 vs G3 = (G7value-G3value)/G3value; ND : Not Determined.

In the present work, L and F lines displayed distinct fillet color and mechanical resistance. Higher fillet lightness associated with higher muscle lipid content is a classical, but not systematically observed, result. This can be due to the higher amount of adipose tissue, a non-pigmented white tissue, located both within muscle tissue and in surrounding connective tissue, especially in myosepta (Lefevre et al., 2015; Marty-Mahe et al., 2004). Moreover, a genetic correlation between fillet lightness and muscle fat content was already observed in other fish species like whitefish (Kause et al., 2011). In the present study, fillets were not pigmented so the main differences in color parameters was measured in yellowness (b*) rather than for redness (a*), and this cannot be attributed to a difference in the retention of carotenoid pigments from food. A similar result was obtained with non-pigmented trout of a similar weight with various muscle fat content resulting from different diet lipid content but only in a low rearing density group (Suarez et al., 2014).

The lower firmness measured for fillet from fish of F line can be related to the higher proportion of adipose tissue, mechanically less resistant than muscle fibers, in fillet. A softer texture associated with a fatter fillet was frequently observed whether higher muscle adiposity results from a dietary origin or from a genetic model of differential adiposity (Johansson et al., 2000; Lefevre et al., 2015; Suarez et al., 2014). Interestingly, with a differential model resulting from more or less lipid in the diet, Suarez et al. (Suarez et al., 2014) showed that the higher firmness of leaner fish was associated with a higher *post-mortem* rigor index. So differential muscle adiposity may also impact the intensity of *post-mortem* events. However, in the present study, higher adiposity in F line was associated with larger white muscle fibers. It has been shown that the size of the muscle fibers has an impact on the salmonid flesh texture, the fillet being firmer the smaller the fibers are (Bugeon et al., 2003; Johnston et al., 2000; Lefevre et al., 2015). So, both higher muscle adiposity and larger fiber may contribute to the observed softer texture in F line.

No difference between the two lines was measured for mechanical resistance of cooked fillet. This confirms the result obtained previously on the third generation of divergent selection (Lefevre et al., 2015) but contradicts a previous study showing that muscle fat content (7.3-14.3 % range) affects the texture of rainbow trout minced cooked fillets (Green-Petersen and Hyldig, 2010). However, the difference measured in mechanical strength of the raw fillets was also found in the smoked fillets. This result seems logical since we applied a traditional cold-smoking process that does not denature muscle proteins. Such a softer smoked fillet texture from fattier fish was previously reported in salmon (Einen et al., 1999; Robb et al., 2002).

Regarding the difference in muscle adiposity generated by divergent selection, the impact on flesh quality may appear limited. Indeed, the difference in Fat-meter^®^ values between the F line and the L line was just over 2 times for the 3^rd^ generation, whereas it was over 4 times for the 7^th^ generation (**Table 6**). We have to keep in mind that fat-meter^®^ evaluates both muscle lipid content and subcutaneous adipose tissue. Interestingly a recent study reports that these two traits were linked to a single region on trout genome suggesting a possibly common mechanism underlying these traits (Blay et al., 2021). However, it should be noted, when comparing the 3^rd^ and 7^th^ generations measured over a long-time interval, that these traits do not only depend on genetics but also on rearing environmental factors (Weil et al., 2013), which can themselves impact the quality parameters, and which were of course different during the rearing of 3^rd^ and 7^th^ generations. For consequences on fillet color parameters, it was difficult to compare the two sets of data as contrary to the third generation, diet was non supplemented with carotenoid for the 7^th^ generation, consequently pigmented and unpigmented fillets were measured respectively for the 3^rd^ and 7^th^ generation of selection. Nevertheless, we could notice that a significant but limited effect of the divergent selection was measured on the color of the fillets both for the 3^rd^ and the 7^th^ generation of selection. So, a greater difference in adiposity did not drastically increase the difference in the color of fillet. For the raw fillet mechanical resistance, when comparing the ratio of the difference between the two lines between the 3^rd^ and the 7^th^ generation of selection (**Table 6**), we observed that this ratio was increased between 2 and 3 folds, depending on the parameter considered. It’s interesting to notice that we measure the same amount of divergence for muscle fatness parameters than for firmness ones. Nevertheless, we observed that the difference between F and L lines in average fibre diameter of the white muscle was increased by more than 10 folds between the 3^rd^ and 7^th^ generation of selection. As mechanical resistance parameters are affected by both fillet adiposity and muscle fibre size, a greater difference in fillet firmness might have been expected. So, we can conclude that the impact, on raw fillet firmness, of muscle adiposity on one hand and those of muscle cellularity on the other hand, do not add up. Raw fillet texture was also shown to be impacted by muscle collagen content or crosslink (Hatae et al., 1986; Li et al., 2005). The muscle connective tissue might also have been impacted by divergent selection on muscle fat content, but this has not been not examined in the present study. Moreover, in another model of differential adiposity in salmon, induced by a more or less energetic diet, and different ration levels, impacting both fillet fat content and firmness, but also fish growth, the hydroxylysyl pyridinoline cross-link concentration was the only factor influencing significantly fillet firmness (Johnsen et al., 2011). The correlated response of adiposity divergent selection on white muscle fiber size makes this biological model more complex to lighten the link between muscle adiposity and flesh quality. However, a negative genetic correlation between muscle fat content and muscle fiber density (−0.76) was also observed in Atlantic salmon, leading to a smaller but significant phenotypic correlation (-0.44), between muscle fat content and fiber density (Vieira et al., 2007). So, the existence of a biological relationship between muscle adiposity and cellularity in salmonid seems likely. Interestingly, higher muscle lipid content induced by high fat diet was shown to be associated to a decrease in the expression of myostatin genes in juvenile rainbow trout (Galt et al., 2014). Even if in fish, myostatin was shown to be not only a specialized strong muscle regulator, but a more general inhibitors of cell proliferation and growth (Gabillard et al., 2013), this suggests a possible relationship between muscle growth regulation and lipid metabolism (Galt et al., 2014). Whatever, whether the lever leading to differential adiposity is genetic or nutritional, it seems relevant to measure the consequences on muscle cellularity to answer this biological question or y to better understand the impacts on product quality.

### 4.2. Some secondary sexual traits were observed in immature fish

Even if all the fishes measured were sexually immature, some significant differences were measured between males and females.

In trout, as in many other fish species, the investment of female in reproduction is more important than male one (Bobe et al., 2010). Indeed, at spawning time the trout GSI is around 15% for female and 7% for male (Bobe et al., 2010). Here, immature 1 year old female had higher GSI values which shows that this trait appears very early in the process of sexual differentiation. We also could notice that, even if interaction between line and sex was not significant, the difference between males and females was more pronounced in F line as fish from this line are more precocious in starting their sexual maturation (Weil et al., 2008). Another dimorphic secondary sexual trait was already present: a larger head for males, compared to females, for both F and L lines. Indeed, in salmonids, sexual maturation is associated with the appearance of a higher development of the lower jaw in males (Monet et al., 2006), and as for GSI, we could notice that this particularly marked phenotype for sexually mature fish seems to appear gradually from the onset of puberty.

We also observed that the diameter of white muscle fibres was higher for males than for females. Such an hypertrophy of muscle fibres in males, before the investment in spermatogenesis, was also observed in Atlantic halibut (Hagen et al., 2006). However, in this work sexual dimorphism was related to a precocious puberty in male, whereas female were immature (Hagen et al., 2006). In our study on trout, both male and female were immature (GSI lower than 1%). And the difference we measured in fibres size was not associated to a difference in fish weight as observed in the study by Hagen and coll. (Hagen et al., 2006). Interestingly, the hyperplasia, which is the recruitment of new muscle fibres, was shown to stop earlier in male than in female in Atlantic halibut (Hagen et al., 2008), which lead to a lower total number of muscle fibres in males, and a different fibre diameter distribution between the two sexes (Hagen et al., 2006). To our knowledge, such a difference in muscle growth pattern, depending on gender, has not been described in salmonids. Nevertheless, the measurement of large salmon (4-5 kg) revealed that males have larger muscle fibres than females (Vieira et al., 2007). In addition, some precocious differences in growth, fillet quality, and fatty acid metabolism between 14 months old rainbow trout males and females were reported (Manor et al., 2015). In this study, no difference between males and females was measured for GSI, but female had higher muscle yield and firmer fillet but muscle cellularity was not measured. The anabolic effect of sex steroids, classically observed, and which leads to the hypertrophy of muscle fibres in males could therefore appear early during gametogenesis in trout and lead to differences as early as pan-size stage. In the present study, difference in fibre size between males and females was essentially observed in F line. This could result from an earlier start of gametogenesis in this line (Weil et al., 2008), resulting in an earlier impact of sexual steroids in male, or to a synergistic effect of muscle lipid content, possibly related to lower myostatin expression (Galt et al., 2014), and sexual steroids in male from F line.

Moreover, the F and L lines divergent genetic model of the present study is a spring-breeding line. Previous studies comparing the evolution of gametogenesis in strains with different spawning seasons have shown that in spring-breeding trout, the onset of gametogenesis starts in the spring, at the same time as for autumn-breeding strain, so one year before the first sexual maturation (Bobe et al., 2010). This may explain the early onset of sexual dimorphism on some traits measured in the present study with pan-size fish.

### 4.3. Relationship between flesh quality parameters

The present model of divergently selected trout is especially relevant to study the relationship between muscle features (adiposity and cellularity) and quality traits without the confounding effect of nutritional factors (different diets or ration levels) or rearing environmental conditions. Moreover, this differential model provides, after seven generations of selection, a wide adiposity range leading to marked differences in muscle traits and quality parameters.

Our data show that some quality parameters, as raw fillet color and mechanical resistance, are highly correlated to both adiposity (fat-meter^®^ value and muscle dry matter content) and muscle cellularity. Some correlations were already observed for the third generation of selection (Lefevre et al., 2015) but the correlation coefficients were lower than those measured in the present study. As fish adiposity was also correlated to muscle cellularity, it’s difficult to conclude whether the difference in quality parameters is attributable to adiposity or muscle fiber size. Indeed, as discussed above, both of them were shown to be determinant of fillet color and texture (Einen et al., 1999; Johnston et al., 2000). Previous work reporting a relationship between fish adiposity and flesh texture have not explored muscle cellularity (Lefevre et al., 2015; Morkore et al., 2001; Robb et al., 2002; Suarez et al., 2014). In contrast, some studies reporting a link between muscle cellularity and fillet texture have measured lipid content (Fauconneau et al., 1993; Johnston et al., 2000; Johnston et al., 2004) and often observe difference in fish adiposity also associated with fillet texture. For example, Fauconneau et al. report that pan size rainbow trout reared à lower temperature (8°C vs 18°C) had both higher lipid content, larger muscle fibers, and less firm raw fillet (Fauconneau et al., 1993). Studying early and late maturing Atlantic salmon strains, Johnston et al. measured a significant positive correlation between muscle cellularity and fillet firmness, whereas muscle cellularity was negatively correlated with lipid content in one strain but not in the other one (Johnston et al., 2000). Still on Atlantic salmon, the comparison of six genetic families submitted to various post-smolt photoperiodic regime showed that fillet from fish maintained with continuous light (24h/day) after sea transfer were firmer, and this was associated with lower lipid content and higher muscle fiber density (Johnston et al., 2004). So, it is notable that in the literature, the link between muscle cellularity and texture is also often associated with different levels of adiposity in salmonids.

In the present work, the PCA analysis confirms that in our model muscle adiposity and cellularity are strongly correlated with each other and negatively correlated with fillet firmness. However, with a multiple linear regression, we can explain 68% of the variability of the raw specific resistance with independent variables: fillet yellowness b*, muscle fibre density, proportion of large muscle fibres (diameter >100µm), and flesh ultimate pH, but the adiposity parameters (fat-meter^®^ value, raw muscle dry matter) were not included in the best model of regression suggesting that, in our model, the main driver of flesh mechanical resistance is muscle fibre size rather than adiposity. The involvement of muscle ultimate pH in this multiple regression model is a bit surprising, since we observe no differences between the two lines for this parameter. However, a significant correlation was obtained between ultimate pH and fillet specific resistance (r=0.39, p<0.01, **Table S2**). Indeed, texture measured at 48h *post-mortem* partly depends on *post-mortem* softening, and ultimate pH can be a good indicator of *post-mortem* evolution processes. Whatever, we must keep in mind that, in some models, textural differences can be attributed to other muscle components than fibers or intra-muscular lipids, especially collagen content or cross-link (Johnston et al., 2006; Li et al., 2005), and that was not measured in the present study.

The consequences of divergent selection on product quality were also measured on smoked fillets. Moreover, smoked fillet data were correlated to those of raw fillet showing similar determinants of quality. Correlations between trout raw and smoked fillet color parameters (lightness, chroma and hue) was also previously reported comparing astaxanthin or canthaxanthin addition in feed (Choubert, 1992). Interestingly, such a relationship between raw and smoked fillet was recently measured in the description of the follow-up of post-spawning trout quality, which is another model of very large variation in fish adiposity (Ahongo et al., 2021). Similarly in the present study, differences in mechanical resistance of smoked fillets can be predicted by those of raw fillets, as it has been observed with post-spawning trout (Ahongo et al., 2021), but not in a previous study measuring large salmon (Birkeland et al., 2004). Nevertheless, we observed that correlation between smoked fillet color and texture and adiposity or muscle cellularity were lower than those obtained for raw fillet. This observation highlights that smoked fillet quality results from the sum of the initial characteristics of the raw product and the impact of the salting / smoking processes.

## 5. Conclusions

The present work confirms that divergent selection on muscle lipid content greatly affects the characteristics of fish and their flesh qualities for both raw, cooked and smoked fillets. Differential adiposity induced by divergent selection affects fillet color and firmness, but the correlated response on muscle cellularity appeared to be the main determinant of fillet firmness. This correlated response to fat-content selection should be examined as a possible biological mechanism linking muscle fat content and muscle growth mechanisms. Further investigations on molecular markers related to muscle characteristics and flesh quality will allow a better understanding of this divergent model.

## 6. CRediT authorship contribution statement

Florence Lefèvre: Conceptualization, Methodology, Investigation, Formal analysis, Writing - Original Draft, Writing - Review & Editing, Visualization, Supervision, Project administration. Jérôme Bugeon: Conceptualization, Methodology, Investigation, Formal analysis, Writing - Review & Editing. Lionel Goardon: Resources, Investigation, Project administration. Thierry Kernéis: Resources, Investigation, Writing - Review & Editing. Laurent Labbé: Resources, Project administration. Stéphane Panserat: Investigation, Writing - Review & Editing. Françoise Médale: Conceptualization. Edwige Quillet: Conceptualization, Methodology, Writing - Review & Editing, Supervision, Project administration.

## 9. Data availability

Data will be made available on request.

## Supporting information

Supplemental tables 1, 2 & 3: correlations between measured traits

## 10. Acknowledgements

The authors thank all the staff of the PEIMA experimental facility (PEIMA, INRAE, Fish Farming systems Experimental Facility, DOI: 10.15454/1.5572329612068406E12, Sizun, France) for rearing the fish over generations of selective breeding and for the present trial and their help for the measurements and sampling at slaughter. They also thank Ms Véronique Lebret and Adeline Jouquan for their technical assistance for the quality measurements.

## References

Ahongo, Y.D., Kerneis, T., Goardon, L., Labbe, L., Bugeon, J., Rescan, P.Y., Lefevre, F., 2021. Flesh quality recovery in female rainbow trout (*Oncorhynchus mykiss*) after spawning. Aquaculture 536. 10.1016/j.aquaculture.2020.736290

Birkeland, S., Rora, A.M.B., Skara, T., Bjerkeng, B., 2004. Effects of cold smoking procedures and raw material characteristics on product yield and quality parameters of cold smoked Atlantic salmon (*Salmo salar* L.) fillets. Food Research International 37(3), 273–286. 10.1016/j.foodres.2003.12.004

Blay, C., Haffray, P., Bugeon, J., D’Ambrosio, J., Dechamp, N., Collewet, G., Enez, F., Petit, V., Cousin, X., Corraze, G., Phocas, F., Dupont-Nivet, M., 2021. Genetic Parameters and Genome-Wide Association Studies of Quality Traits Characterised Using Imaging Technologies in Rainbow Trout, *Oncorhynchus mykiss*. Front. Genet. 12, 17. 10.3389/fgene.2021.639223

Bobe, J., Breton, B., Fostier, A., Guiguen, Y., Jalabert, B., Kah, O., Labbé, C., Lareyre, J.J., Le Bail, P.Y., Le Gac, F., Leveroni Calvi, S., Mahé, S., Quillet, E., Vandeputte, M., 2010. Sexualité et reproduction, in: QUAE (Ed.), La truite arc-en-ciel. De la biologie à l’élevage. pp. 39–81.

Bugeon, J., Lefevre, F., Fauconneau, B., 2003. Fillet texture and muscle structure in brown trout (*Salmo trutta*) subjected to long-term exercise. Aquaculture Research 34(14), 1287–1295. 10.1046/j.1365-2109.2003.00938.x

Choubert, G., 1992. Salmonid pigmentation: Dynamics and factors of variation. A review. INRA Productions Animales 5(4), 235–246. 10.20870/productions-animales.1992.5.4.4237

CIE, 1976. CIE, Commission Internationale de l’Éclairage, Colorimetry, Publication no 15, Bureau central de la CIE, Vienna, Austria (1976) 14 pp.

Douirin, C., Haffray, P., Vallet, J.L., Fauconneau, B., 1998. Determination of the lipid content of rainbow trout (*Oncorhynchus mykiss*) fillets with the Torry Fish Fat Meter^(R)^. Sciences des aliments 18(5), 527–535.

Einen, O., Morkore, T., Rora, A.M.B., Thomassen, M.S., 1999. Feed ration prior to slaughter - a potential tool for managing product quality of Atlantic salmon (*Salmo salar*). Aquaculture 178(1-2), 149–169. 10.1016/S0044-8486(99)00126-X

Fauconneau, B., Chmaitilly, J., Andre, S., Cardinal, M., Cornet, J., Vallet, J.L., Dumont, J.P., Laroche, M., 1993. Characteristics of rainbow trout flesh.2. Physical and sensory aspects. Sciences des aliments 13(2), 189–199.

Gabillard, J.C., Biga, P.R., Rescan, P.Y., Seiliez, I., 2013. Revisiting the paradigm of myostatin in vertebrates: Insights from fishes. General and Comparative Endocrinology 194, 45–54. 10.1016/j.ygcen.2013.08.012

Galt, N.J., Froehlich, J.M., Meyer, B.M., Barrows, F.T., Biga, P.R., 2014. High-fat diet reduces local myostatin-1 paralog expression and alters skeletal muscle lipid content in rainbow trout, *Oncorhynchus mykiss*. Fish Physiology and Biochemistry 40(3), 875–886. 10.1007/s10695-013-9893-4

Golik, W., Dameron, O., Bugeon, J., Fatet, A., Hue, I., Hurtaud, C., Reichstadt, M., Meunier-Salaün, M.C., Vernet, J., Joret, L., Papazian, F., Nédellec, C., Le Bail, P.Y., 2012. ATOL: the multi-species livestock trait ontology. 6th International Conference on Metadata and Semantic Research 28-30 Nov.2012, Cadiz, Spain.

Green-Petersen, D.M.B., Hyldig, G., 2010. Variation in sensory profile of individual rainbow trout (*Oncorhynchus mykiss*) from the same production batch. Journal of Food Science 75(9), S499–S505. 10.1111/j.1750-3841.2010.01830.x

Haffray, P., Bugeon, J., Pincent, C., Chapuis, H., Mazeiraud, E., Rossignol, M.N., Chatain, B., Vandeputte, M., Dupont-Nivet, M., 2012. Negative genetic correlations between production traits and head or bony tissues in large all-female rainbow trout (*Oncorhynchus mykiss*). Aquaculture 368, 145–152. 10.1016/j.aquaculture.2012.09.023

Hagen, O., Solberg, C., Johnston, I.A., 2006. Sexual dimorphism of fast muscle fibre recruitment in farmed Atlantic halibut (*Hippoglossus hippoglossus* L.). Aquaculture 261(4), 1222–1229. 10.1016/j.aquaculture.2006.09.026

Hagen, O., Vieira, V.L.A., Solberg, C., Johnston, I.A., 2008. Myotube production in fast myotomal muscle is switched-off at shorter body lengths in male than female Atlantic halibut *Hippoglossus hippoglossus* (L.) resulting in a lower final fibre number. Journal of Fish Biology 73(1), 139–152. 10.1111/j.1095-8649.2008.01917.x

Hatae, K., Tobimatsu, A., Takeyama, M., Matsumoto, J.J., 1986. Contribution of connective tissues on the texture difference of various fish species. Nippon Suisan Gakkaishi 52(11), 2001–2007.

Johansson, L., Kiessling, A., Kiessling, K.H., Berglund, L., 2000. Effects of altered ration levels on sensory characteristics, lipid content and fatty acid composition of rainbow trout (*Oncorhynchus mykiss*). Food Quality and Preference 11(3), 247–254. 10.1016/S0950-3293(99)00073-7

Johnsen, C.A., Hagen, O., Adler, M., Jonsson, E., Kling, P., Bickerdike, R., Solberg, C., Bjornsson, B.T., Bendiksen, E.A., 2011. Effects of feed, feeding regime and growth rate on flesh quality, connective tissue and plasma hormones in farmed Atlantic salmon (*Salmo salar* L.). Aquaculture 318(3-4), 343–354. 10.1016/j.aquaculture.2011.05.040

Johnston, I.A., Alderson, R., Sandham, C., Dingwall, A., Mitchell, D., Selkirk, C., Nickell, D., Baker, R., Robertson, B., Whyte, D., Springate, J., 2000. Muscle fibre density in relation to the colour and texture of smoked Atlantic salmon (*Salmo salar* L.). Aquaculture 189(3-4), 335–349. 10.1016/S0044-8486(00)00373-2

Johnston, I.A., Li, X.J., Vieira, V.L.A., Nickell, D., Dingwall, A., Alderson, R., Campbell, P., Bickerdike, R., 2006. Muscle and flesh quality traits in wild and farmed Atlantic salmon. Aquaculture 256(1-4), 323–336. 10.1016/j.aquaculture.2006.02.048

Johnston, I.A., Manthri, S., Bickerdike, R., Dingwall, A., Luijkx, R., Campbell, P., Nickell, D., Alderson, R., 2004. Growth performance, muscle structure and flesh quality in out-of-season Atlantic salmon (*Salmo salar*) smolts reared under two different photoperiod regimes. Aquaculture 237(1-4), 281–300. 10.1016/j.aquaculture.2004.04.026

Kamalam, B.S., Medale, F., Kaushik, S., Polakof, S., Skiba-Cassy, S., Panserat, S., 2012. Regulation of metabolism by dietary carbohydrates in two lines of rainbow trout divergently selected for muscle fat content. Journal of Experimental Biology 215(15), 2567–2578. 10.1242/jeb.070581

Kause, A., Quinton, C., Airaksinen, S., Ruohonen, K., Koskela, J., 2011. Quality and production trait genetics of farmed European whitefish, *Coregonus lavaretus*. Journal of Animal Science 89(4), 959–971. 10.2527/jas.2010-2981

Kolditz, C., Borthaire, M., Richard, N., Corraze, G., Panserat, S., Vachot, C., Lefevre, F., Medale, F., 2008. Liver and muscle metabolic changes induced by dietary energy content and genetic selection in rainbow trout (*Oncorhynchus mykiss*). American Journal of Physiology - Regulatory Integrative and Comparative Physiology 294(4), R1154–R1164. 10.1152/ajpregu.00766.2007

Lefevre, F., Cardinal, M., Bugeon, J., Labbe, L., Medale, F., Quillet, E., 2015. Selection for muscle fat content and triploidy affect flesh quality in pan-size rainbow trout, *Oncorhynchus mykiss*. Aquaculture 448, 569–577. 10.1016/j.aquaculture.2015.06.029

Li, X.J., Bickerdike, R., Lindsay, E., Campbell, P., Nickell, D., Dingwall, A., Johnston, I.A., 2005. Hydroxylysyl pyridinoline cross-link concentration affects the textural properties of fresh and smoked Atlantic salmon (*Salmo salar* L.) flesh. Journal of Agricultural and Food Chemistry 53(17), 6844–6850. 10.1021/jf050743+

Lopez-De Leon, A., Rojkind, M., 1985. A simple micromethod for collagen and total protein determination in formalin-fixed paraffin-embedded sections. Journal of Histochemistry & Cytochemistry 33(8), 737–743. 10.1177/33.8.2410480

Lutfi, E., Gong, N.P., Johansson, M., Sanchez-Moya, A., Bjornsson, B.T., Gutierrez, J., Navarro, I., Capilla, E., 2018. Breeding selection of rainbow trout for high or low muscle adiposity differentially affects lipogenic capacity and lipid mobilization strategies to cope with food deprivation. Aquaculture 495, 161–171. 10.1016/j.aquaculture.2018.05.039

Manor, M.L., Cleveland, B.M., Kenney, P.B., Yao, J.B., Leeds, T., 2015. Differences in growth, fillet quality, and fatty acid metabolism-related gene expression between juvenile male and female rainbow trout. Fish Physiology and Biochemistry 41(2), 533–547. 10.1007/s10695-015-0027-z

Marty-Mahe, P., Loisel, P., Fauconneau, B., Haffray, P., Brossard, D., Davenel, A., 2004. Quality traits of brown trouts (*Salmo trutta*) cutlets described by automated color image analysis. Aquaculture 232(1-4), 225–240. 10.1016/S0044-8486(03)00458-7

Medale, F., 2009. Lipid content and fatty acid composition of the flesh of fish from fisheries and farming. Cahiers de Nutrition et de Dietetique 44(4), 173–181. 10.1016/j.cnd.2009.04.002

Monet, G., Uyanik, A., Champigneulle, A., 2006. Geometric morphometrics reveals sexual and genotypic dimorphisms in the brown trout. Aquatic Living Resources 19(1), 47–57. 10.1051/alr:2006004

Morkore, T., Vallet, J.L., Cardinal, M., Gomezguillen, M.C., Montero, R., Torrissen, O.J., Nortvedt, R., Sigurgisladottir, S., Thomassen, M.S., 2001. Fat content and fillet shape of Atlantic salmon: Relevance for processing yield and quality of raw and smoked products. Journal of Food Science 66(9), 1348–1354. 10.1111/j.1365-2621.2001.tb15213.x

Quillet, E., Le Guillou, S., Aubin, J., Fauconneau, B., 2005. Two-way selection for muscle lipid content in pan-size rainbow trout (*Oncorhynchus mykiss*). Aquaculture 245(1-4), 49–61. 10.1016/j.aquaculture.2004.12.014

Quillet, E., Le Guillou, S., Aubin, J., Labbe, L., Fauconneau, B., Medale, F., 2007. Response of a lean muscle and a fat muscle rainbow trout (*Oncorhynchus mykiss*) line on growth, nutrient utilization, body composition and carcass traits when fed two different diets. Aquaculture 269(1-4), 220–231. 10.1016/j.aquaculture.2007.02.047

Robb, D.H.F., Kestin, S.C., Warriss, P.D., Nute, G.R., 2002. Muscle lipid content determines the eating quality of smoked and cooked Atlantic salmon (*Salmo salar*). Aquaculture 205(3-4), 345–358. 10.1016/S0044-8486(01)00710-4

Suarez, M.D., Garcia-Gallego, M., Trenzado, C.E., Guil-Guerrero, J.L., Furne, M., Domezaine, A., Alba, I., Sanz, A., 2014. Influence of dietary lipids and culture density on rainbow trout (*Oncorhynchus mykiss*) flesh composition and quality parameter. Aquacultural Engineering 63, 16–24. 10.1016/j.aquaeng.2014.09.001

Szczesniak, A.S., Humbaugh, P.R., Block, H.W., 1970. Behavior of different foods in the standard shear compression cell of the shear press and the effect of sample weigth on peak area and maximum force. Journal of Texture Studies 1, 356–378. 10.1111/j.1745-4603.1970.tb00736.x

Tobin, D., Kause, A., Mantysaari, E.A., Martin, S.A.M., Houlihan, D.F., Dobly, A., Kiessling, A., Rungruangsak-Torrissen, K., Ritola, O., Ruohonen, K., 2006. Fat or lean? The quantitative genetic basis for selection strategies of muscle and body composition traits in breeding schemes of rainbow trout (*Oncorhynchus mykiss*). Aquaculture 261(2), 510–521. 10.1016/j.aquaculture.2006.07.023

Vieira, V.L.A., Norris, A., Johnston, I.A., 2007. Heritability of fibre number and size parameters and their genetic relationship to flesh quality traits in Atlantic salmon (*Salmo salar* L.). Aquaculture 272, S100–S109. 10.1016/j.aquaculture.2007.08.028

Weil, C., Goupil, A.S., Quillet, E., Labbe, L., Le Gac, F., 2008. Two-way selection for muscle lipid content modifies puberty and gametogenesis in rainbow trout. Cybium 32(2), 198–198.

Weil, C., Lefevre, F., Bugeon, J., 2013. Characteristics and metabolism of different adipose tissues in fish. Reviews in Fish Biology and Fisheries 23(2), 157–173. 10.1007/s11160-012-9288-0

