## Supplemental tables 1, 2 & 3: correlations between measured traits for "From the third to the seventh generation of selection for muscle fat content in rainbow trout: Consequences for flesh quality"

(1) : INRAE, LPGP, 35000, Rennes, France

(2) : INRAE, PEIMA, 29450 Sizun, France

(3) : INRAE, Univ. Pau & Pays Adour, NUMEA, 64310, Saint-Pée-sur-Nivelle, France

(4) : Université Paris-Saclay, INRAE, AgroParisTech, GABI, 78350 Jouy-en-Josas, France

* Correspondence to:
Florence LEFÈVRE
INRAE,
Fish Physiology and Genomics Institute (LPGP),
Campus de Beaulieu,
35042 Rennes cedex,
France


### Supplementary tables

**Table S1**: Pearson correlation within raw and cooked fillet color parameters and between color parameters and others measured parameters, n=60.

| | BW | K | Fat | FD | FM D | % SF | % LF | pHi | L*s | a*s | b*s | dmr | pHu | L*r | a*r | b*r | dmc | pHc |
|---|---|---|---|---|---|---|---|---|---|---|---|---|---|---|---|---|---|---|
| **pHi** | 0.16 NS | 0.19 NS | 0.09 NS | -0.16 NS | 0.21 NS | -0.24 NS | 0.17 NS | - | | | | | | | | | | |
| **L*s** | 0.32 * | 0.46 *** | 0.57 *** | -0.62 *** | 0.65 *** | -0.59 *** | 0.61 *** | 0.09 NS | - | | | | | | | | | |
| **a*s** | -0.05 NS | 0.21 NS | 0.31 * | -0.09 NS | 0.19 NS | -0.13 NS | 0.19 NS | 0.26 * | 0.17 NS | - | | | | | | | | |
| **b*s** | 0.20 NS | 0.62 *** | 0.60 *** | -0.48 *** | 0.53 *** | -0.51 *** | 0.50 *** | 0.35 ** | 0.67 *** | 0.60 *** | - | | | | | | | |
| **dmr** | 0.02 NS | 0.73 *** | 0.94 *** | -0.76 *** | 0.77 *** | -0.73 *** | 0.76 *** | 0.14 NS | 0.56 *** | 0.34 ** | 0.64 *** | - | | | | | | |
| **pHu** | -0.44 *** | -0.21 NS | -0.19 NS | 0.19 NS | -0.24 NS | 0.15 NS | -0.24 NS | 0.20 NS | -0.55 *** | -0.09 NS | -0.29 * | -0.23 NS | - | | | | | |
| **L*r** | 0.22 NS | 0.49 NS | 0.63 *** | -0.59 *** | 0.65 *** | -0.64 *** | 0.59 *** | 0.28 * | 0.74 *** | 0.29 * | 0.70 *** | 0.61 *** | -0.32 * | - | | | | |
| **a*r** | -0.14 NS | 0.1 NS | 0.33 * | -0.07 NS | 0.16 NS | -0.08 NS | 0.17 NS | 0.04 NS | 0.14 NS | 0.72 *** | 0.34 ** | 0.33 * | -0.20 NS | 0.04 NS | - | | | |
| **b*r** | 0.09 NS | 0.51 *** | 0.71 *** | -0.59 *** | 0.60 *** | -0.59 *** | 0.59 *** | -0.08 NS | 0.79 *** | 0.27 * | 0.69 *** | 0.69 *** | -0.56 *** | 0.59 *** | 0.38 ** | - | | |
| **dmc** | -0.08 NS | 0.62 *** | 0.89 *** | -0.69 *** | 0.71 *** | -0.73 *** | 0.64 *** | 0.13 NS | 0.59 *** | 0.32 * | 0.66 *** | 0.89 *** | -0.21 NS | 0.67 *** | 0.30 * | 0.70 *** | - | |
| **pHc** | -0.03 NS | 0.04 NS | -0.08 NS | -0.00 NS | 0.04 NS | 0.07 NS | 0.03 NS | -0.25 NS | -0.08 NS | -0.21 NS | -0.19 NS | -0.11 NS | 0.39 ** | -0.22 NS | 0.27 * | -0.14 NS | -0.17 NS | - |
| **L*c** | 0.47 *** | 0.09 NS | -0.14 NS | -0.09 NS | 0.04 NS | -0.04 NS | 0.09 NS | -0.13 NS | 0.16 NS | -0.18 NS | -0.02 NS | -0.08 NS | -0.42 ** | -0.06 NS | -0.03 NS | 0.16 NS | -0.12 NS | -0.08 NS |
| **a*c** | 0.42 ** | 0.42 ** | 0.19 NS | -0.19 NS | 0.19 NS | -0.22 NS | 0.21 NS | 0.33 * | 0.22 NS | 0.34 ** | 0.46 *** | 0.26 * | -0.31 * | 0.23 NS | 0.26 * | 0.24 NS | 0.19 NS | -0.29 * |
| **b*c** | 0.26 * | 0.47 *** | 0.43 ** | -0.34 ** | 0.38 ** | -0.35 ** | 0.34 ** | 0.21 NS | 0.38 ** | 0.25 NS | 0.60 *** | 0.47 *** | -0.37 ** | 0.43 ** | 0.25 NS | 0.50 *** | 0.44 ** | -0.29 * |

BW: body weight; K: condition factor; Fat: Fat-meter value; FD: Fiber density; FMD: Fiber Mean Diameter; % SF: % of Small Fibers; %LF: % of Large Fibers; pHi: initial pH; L*s, a*s, b*s: lightness, redness, yellowness at slaughter; dmr: raw muscle dry matter content; pHu: ultimate pH; L*r, a*r, b*r: lightness, redness, yellowness for raw flesh at 48h *post-mortem*; dmc: cooked muscle dry matter content; pHc: cooked pH; L*c, a*c, b*c: lightness, redness, yellowness for cooked flesh; NS: p≥0.05, *: p<0.05, **: p<0.01, ***: p<0.001.

**<u>Table S2</u>**: Pearson correlation within raw and cooked fillet mechanical resistance parameters and between mechanical resistance parameters and others measured parameters, n=60.

| | **PT-MF-s** | **PT-W/wi-s** | **PT-MF-r** | **PT-W/wi-r** | **KSP-MF-r** | **KSP-MF/we-r** | **PT-MF-c** | **PT-W/wi-c** | **KSP-MF-c** | **KSP-MF/we-c** |
|---|---|---|---|---|---|---|---|---|---|---|
| **PT-W/wi-s** | 0.61 *** | - | | | | | | | | |
| **PT-MF-r** | 0.38 ** | 0.53 *** | - | | | | | | | |
| **PT-W/wi-r** | 0.01 NS | 0.35 ** | 0.76 *** | - | | | | | | |
| **KSP-MF-r** | 0.40 ** | 0.56 *** | 0.65 *** | 0.47 *** | - | | | | | |
| **KSP-MF/we-r** | 0.56 *** | 0.35 ** | 0.32 * | -0.03 NS | -0.34 ** | - | | | | |
| **PT-MF-c** | -0.08 NS | 0.14 NS | 0.20 NS | 0.24 NS | 0.31 * | -0.26 * | - | | | |
| **PT-W/wi-c** | 0.15 NS | 0.28 * | 0.28 * | 0.17 NS | 0.28 * | 0.01 NS | 0.83 *** | - | | |
| **KSP-MF-c** | -0.20 NS | -0.02 NS | 0.05 NS | 0.21 NS | 0.36 ** | -0.34 ** | 0.30 * | 0.06 NS | - | |
| **KSP-MF/we-c** | 0.05 NS | -0.17 NS | -0.27 * | -0.36 ** | -0.21 NS | 0.33 * | -0.18 NS | -0.04 NS | 0.14 NS | - |
| **BW** | -0.04 NS | 0.14 NS | 0.32 * | 0.45 *** | 0.60 *** | -0.37 ** | 0.41 ** | 0.15 NS | 0.68 *** | -0.44 *** |
| **K** | -0.44 *** | -0.21 NS | 0.03 NS | 0.29 * | -0.13 NS | -0.57 *** | 0.15 NS | -0.15 NS | 0.24 NS | -0.53 *** |
| **Fat** | -0.54 *** | -0.37 ** | -0.33 * | 0.11 NS | -0.56 *** | -0.62 *** | -0.05 NS | -0.24 NS | 0.01 NS | -0.23 NS |
| **FD** | 0.41 ** | 0.21 NS | 0.15 NS | -0.28 * | 0.33 * | 0.73 *** | -0.15 NS | 0.07 NS | -0.19 NS | 0.36 ** |
| **FMD** | -0.42 ** | -0.25 NS | -0.19 NS | 0.26 * | -0.32 * | -0.72 *** | 0.14 NS | -0.08 NS | 0.22 NS | -0.33 * |
| **% SF** | 0.56 *** | 0.30 * | 0.24 NS | -0.16 NS | 0.35 ** | 0.69 *** | -0.09 NS | 0.13 NS | -0.17 NS | 0.32 * |
| **% LF** | -0.35 ** | -0.21 NS | -0.15 NS | 0.32 * | -0.26 * | -0.66 *** | 0.13 NS | -0.09 NS | 0.21 NS | -0.33 * |
| **pHi** | 0.13 NS | -0.01 NS | -0.22 NS | -0.18 NS | 0.01 NS | -0.14 NS | -0.03 NS | -0.16 NS | -0.01 NS | -0.37 ** |
| **dmr** | -0.53 *** | -0.37 ** | -0.26 * | -0.18 NS | -0.50 *** | -0.66 *** | 0.01 NS | -0.24 NS | 0.11 NS | -0.29 * |
| **pHu** | 0.23 NS | -0.01 NS | 0.03 NS | -0.20 NS | -0.14 NS | 0.39 ** | -0.26 * | -0.08 NS | -0.54 *** | -0.06 NS |
| **dmc** | -0.57 *** | -0.43 ** | -0.39 ** | -0.04 NS | -0.58 *** | -0.66 *** | -0.10 NS | -0.27 * | 0.05 NS | -0.24 NS |
| **pHc** | 0.15 NS | 0.19 NS | 0.39 ** | 0.40 ** | 0.13 NS | 0.19 NS | 0.00 NS | 0.18 NS | -0.29 * | -0.29 * |

PT : Puncture Test; KSP : Kramer Shear Press, MF : maximal force, wi : sample width; we : sample weight; 's' : measurement at slaughter; 'r' : raw fillet measured at 48h *post-mortem*; 'c' : cooked fillet; BW: body weight; K: condition factor; Fat: Fat-meter value; FD: Fiber density; FMD: Fiber Mean Diameter; % SF: % of Small Fibers; %LF: % of Large Fibers; pHi: initial pH; dmr: raw muscle dry matter content; pHu: ultimate pH; dmc: cooked muscle dry matter content; pHc: cooked pH; NS: $p \geq 0.05$, *: $p<0.05$, **: $p<0.01$, ***: $p<0.001$.

**Table S3**: Pearson correlation within smoked fillet instrumentally measured quality parameters variables, and between smoked fillet quality parameters variables and others measured parameters, n=60.

| | dm-smok | pH-smok | L*-smok | a*-smok | b*-smok | PT-MF-smok | PT-W/wi-smok | KSP-MF-smok | KSP-MF/we--smok |
|---|---|---|---|---|---|---|---|---|---|
| pH-smok | -0.04 NS | - | | | | | | | |
| L*-smok | 0.03 NS | -0.36 ** | - | | | | | | |
| a*-smok | 0.59 *** | 0.21 NS | -0.10 NS | - | | | | | |
| b*-smok | 0.60 *** | -0.02 NS | -0.14 NS | 0.33 * | - | | | | |
| PT-MF-smok | -0.35 ** | 0.08 NS | 0.04 NS | -0.34 ** | -0.28 * | - | | | |
| PT-W/wi-smok | 0.02 NS | 0.02 NS | 0.08 NS | -0.13 NS | 0.15 NS | 0.75 *** | - | | |
| KSP-MF-smok | -0.21 NS | -0.07 NS | -0.31 * | -0.40 ** | 0.02 NS | 0.29 * | 0.36 ** | - | |
| KSP-MF/we--smok | -0.17 NS | 0.15 NS | -0.34 ** | -0.43 ** | 0.04 NS | 0.57 *** | 0.52 *** | 0.63 *** | - |
| BW | -0.20 NS | -0.23 NS | 0.20 NS | 0.06 NS | -0.14 NS | -0.26 * | -0.18 NS | 0.20 NS | -0.54 *** |
| K | 0.38 ** | 0.05 NS | 0.24 NS | 0.30 * | -0.03 NS | -0.29 * | -0.17 NS | -0.35 ** | -0.44 ** |
| Fat | 0.92 *** | 0.06 NS | -0.02 NS | 0.59 *** | 0.59 *** | -0.33 ** | -0.02 NS | -0.31 * | -0.21 NS |
| pHi | 0.09 NS | 0.06 NS | 0.04 NS | 0.21 NS | -0.11 NS | 0.05 NS | -0.06 NS | -0.35 ** | -0.10 NS |
| L*s | 0.41 ** | 0.01 NS | 0.25 NS | 0.30 * | 0.33 * | -0.35 ** | 0.00 NS | -0.13 NS | -0.47 *** |
| a*s | 0.43 ** | -0.33 * | 0.32 * | 0.44 *** | 0.30 * | -0.00 NS | 0.09 NS | -0.21 NS | -0.12 NS |
| b*s | 0.66 *** | -0.38 ** | 0.31 * | 0.42 ** | 0.35 ** | -0.08 NS | 0.20 NS | -0.13 NS | -0.21 NS |
| PT-MF-s | -0.23 NS | 0.12 NS | -0.00 NS | -0.27 NS | -0.19 NS | 0.69 *** | 0.49 *** | 0.32 * | 0.42 ** |
| PT-W/wi-s | -0.21 NS | 0.05 NS | 0.02 NS | -0.15 NS | -0.03 NS | 0.56 *** | 0.54 *** | 0.34 ** | 0.34 ** |

dm-smok: smoked fillet dry matter content; pH-smok: smoked fillet ultimate pH; L*-smok, a*-smok, b*-smok: smoked fillet lightness, redness, yellowness; PT : Puncture Test; KSP : Kramer Shear Press, MF : maximal force, wi : sample width; we : sample weight; 'smok' : smoked fillet measured; BW: body weight; K: condition factor; Fat: Fat-meter value; pHi: initial pH; L*s, a*s, b*s: lightness, redness, yellowness at slaughter; 's' : measurement at slaughter; NS: p≥0.05, *: p<0.05, **: p<0.01, ***: p<0.001.
